# Microbiome-derived hydroxyphenyl propanoates enhance antitumour immunity by potentiating gasdermin D activity in tumour-associated myeloid cells

**DOI:** 10.64898/2026.04.23.720410

**Authors:** Saif Sikdar, Dae-Sun Kim, Tie Wang, Sacha Benaoudia, Jahanara Rajwani, Kristofor K Ellestad, Pauline Douglas, Ruzena Filandrova, Ted Verhey, Katrina I Ellestad, Marija Drikic, Darren Derksen, Ian Lewis, Marco Gallo, Sorana Morrissy, David Schriemer, Kathy McCoy, Franz J. Zemp, Douglas J. Mahoney

**Affiliations:** Arnie Charbonneau Cancer Institute, University of Calgary; Calgary, AB, Canada; Alberta Children’s Hospital Research Institute, University of Calgary, Calgary, AB, Canada; Department of Microbiology, Immunology, and Infectious Diseases, Cumming School of Medicine, University of Calgary, Calgary, AB, Canada; The Calvin, Joan and Phoebe Snyder Institute for Chronic Disease, University of Calgary, Calgary, AB, Canada; Department of Biochemistry and Molecular Biology, Cumming School of Medicine, University of Calgary; Calgary, AB, Canada; Institute of Organic Chemistry and Biochemistry of the Czech Academy of Sciences, Prague, Czech Republic; Department of Biological Sciences, University of Calgary, Calgary, AB, Canada; Department of Chemistry, University of Calgary, Calgary, AB, Canada; Department of Physiology and Pharmacology, Cumming School of Medicine, University of Calgary, Calgary, AB, Canada

**Keywords:** Cancer, Immunotherapy, Microbiome, Metabolomics, scRNAseq, Thermal proteome profiling, Gasdermin D, Immunomodulation, Hydroxyphenyl propanoates, Immune checkpoint blockade

## Abstract

The microbiome plays a critical role in immune health and response to immune-stimulating treatments, including cancer immunotherapy. However, the molecular mechanisms by which commensal microbes influence cancer immunology remain poorly understood. Here, we report a class of microbiome-derived metabolites called hydroxyphenyl propanoates (HPP) that enhance tumour immune surveillance and response to immune checkpoint blockade (ICB) therapy in mice. HPPs function as potentiators of innate immune signalling in tumour-associated myeloid cells by promoting cleavage of the pore-forming protein gasdermin D (GSDMD), a critical effector of inflammasome function. This enhances the secretion of proinflammatory cytokines, such as IL-1 family members, while simultaneously protecting against pyroptosis. Secreted IL-1 family cytokines in turn elicit autocrine and paracrine inflammatory signaling in tumour-infiltrating leukocytes. HPP supplementation increases anticancer T cell function, improves disease control and extends overall survival (OS) in tumour-bearing mice harboring complex microbiomes or those treated with broad-spectrum antibiotics. In humans, GSDMD activation is associated with a favourable response to PD-1 blockade therapy in patients with advanced stage melanoma. Our study uncovers a molecular mechanism of regulation over host antitumour immunity that is modifiable and can be harnessed for improved cancer immunotherapy in a broad spectrum of patients.

## INTRODUCTION

Successful cancer immune surveillance and immunotherapy depends on dynamic interactions between tumours and cells of the innate and adaptive immune system^1,2^. These interactions are complex and influenced by the cancer’s mutations, the host’s genome and epigenome, and extrinsic factors such as the host’s exposome, including past treatments^3–5^. As the molecular mechanisms underpinning cancer-immune interactions have been unraveled^6,7^, powerful new therapeutic opportunities have been uncovered. These include ICB therapy to relieve anticancer T cells from immunosuppressive signaling in the tumour microenvironment (TME)^8^ and other highly effective immune-based treatments such as adoptive and engager T cell therapeutics^9^.

Over the past decade, the role played by commensal microbes in cancer-immune interactions has gained considerable attention^10–18^. Studies have shown that an intact microbiome is required for a strong immune response to cancer, with germ-free mice being hypo-responsive to cancer immunotherapy and broad-spectrum antibiotic treatment being generally detrimental to anticancer immunity in mice and humans^13,19–21^. Deep sequencing of fecal samples has identified *Bacteroides, Bifidobacterium* and *Enterococcus*, amongst other bacterial genera, as being positively associated with immunotherapy responses in mice and humans^11–13,22,23^. In line with this, fecal transplant from ICB-responsive or non-responsive patients confers matched responsiveness or non-responsiveness to ICB treatment in mice^13–15^. Moreover, reconstitution of individual microbes associated with a positive response improves immunotherapy outcomes in antibiotic-treated or germ-free mice^11,13^. Collectively, these findings have prompted several clinical trials evaluating fecal transplantation neoadjuvant to ICB therapy, which have yielded promising early results^23,24^. However, fecal transplant is a crude therapeutic that is not easily scalable and, in addition to its bioactive microbes, contains many additional and potentially harmful components. Deeper knowledge of microbiome-immune interactome should enable the development of tailored, more sophisticated microbial therapeutics with greater potency in patients, such as individual microbes, defined consortia of microbes, or bacteria genetically modified for enhanced anticancer immunity^18,22,25^.

While promising, microbes are complex therapeutics that are cumbersome and expensive to manufacture, requiring highly specialized current good manufacturing practice (cGMP) facilities, and are subject to a high degree of regulatory scrutiny^26^. Given similar challenges faced by mammalian cell therapies, and the difficulties that has posed even when they are highly effective^26,27^, a simpler approach to leveraging microbe-host interactions to boost anticancer immunity is desirable. Moreover, host resistance to colonization and susceptibility to broad spectrum antibiotics represents difficult challenges facing live bacterial therapeutics in cancer patients. As commensal microbes commonly interact with host immune cells via secreted metabolites^28,29^, a different approach is to develop those bioactive molecules into natural or drug products. Indeed, recent studies have found that the SCFAs pentanoate, butyrate, and propionate, amongst others, as well as other microbe-derived metabolites such as Trimethylamine N-Oxide (TMAO), inosine, indole-3-proprionic acid, desaminotyrosine and tryptophan regulate immune responses to cancer through a variety of molecular mechanisms^16,30–35^. Microbiome-derived metabolites are attractive therapeutic platforms, as they are natural to humans, selected for bioactivity *in vivo*, and have been evolved to function in the complex biological milieu of host-tumour-microbe interactions. Moreover, delivery of rationally selected microbiome-derived metabolites allows for the isolation of a specific bioactivity from a complex microbial community, mitigating potential toxicities that could be derived from other bioactivities in that ecosystem. However, the relationship between the microbiome-derived metabolome and cancer immune surveillance and immunotherapy remains relatively poorly understood. The small handful of metabolites known to promote anticancer immunity likely do not capture the full breadth of interactions between molecules secreted from the microbiome, cancer and the immune system. A deeper understanding of beneficial metabolites and their interactions with cancer-associated immune compartments could guide the development of novel immune-boosting therapeutics, independent of a patient’s microbiome composition or antibiotic exposure.

In this study, we conducted multi-omics profiling of genetically identical mice harbouring diverse microbiomes to elucidate interactions between commensal microbe-derived metabolites, solid tumours, and the host immune system. We identify HPPs as a class of microbiome-derived metabolite that improves cancer immune surveillance and response to ICB therapy in mice by potentiating proinflammatory cytokine secretion through GSDMD pores in tumour-infiltrating myeloid cells. HPPs also work in human cells, and their bioactivity is associated with a better response to ICB therapy in melanoma patients. This mechanism may help explain divergent responses to immunotherapy in human patients and represents a potential neoadjuvant treatment for improving immunotherapy outcomes in cancers rich in immunosuppressive myeloid cells, even in patients harboring disparate microbiomes or undergoing treatment with broad-spectrum antibiotics.

## RESULTS

### The microbiome impacts tumour immune surveillance in genetically identical hosts

Studies have shown that inbred C57BL/6 mice obtained from different laboratories harbor divergent microbiomes and have differing capacities for cancer immune surveillance^11,36^. To explore the relationship between complex microbiomes and anticancer immunity, we first sought to confirm these observations in mice obtained from Charles River Laboratories (CR), Taconic Biosciences (TAC) and The Jackson Laboratory (JAX). As expected, 16S sequencing of fecal samples showed that C57BL/6 mice obtained from different repositories were colonized with vastly divergent microbiomes (Extended figure 1a, b). Consistent with previous studies^11,36^, *Muribaculaceae*, previously known as *Bacteroidales S24-7*, was the most abundant fecal bacterial family in JAX mice. In contrast, *Bacteroidaceae* was the predominant fecal bacterial family in CR and TAC mice. To test if these divergent microbiomes affected tumour growth or cancer immune surveillance, we orthotopically implanted M3-9-M rhabdomyosarcoma (RMS) cells bearing H-Y histocompatibility antigens^37^ into female mice obtained from CR, TAC or JAX. OS was extended in JAX compared to CR or TAC mice (Extended figure 1c-f), an effect that was dependent on CD8^+^ T cells (Extended figure 1g-h). In a complementary approach, we used orally delivered broad-spectrum antibiotics (ABX) to reduce fecal bacterial abundance (Extended figure 1i,j) and found that ABX treatment extended survival in CR mice but reduced survival in JAX mice (Extended figure 1k,l). Collectively, these data are consistent with previous studies suggesting that CR and TAC mice are colonized with microbes that suppress anticancer immunity whereas JAX mice are colonized with microbes that support anticancer immunity^11,36^.

However, there are two important limitations to these experimental approaches: 1.) C57BL/6 mice sourced from different repositories are genetically distinct because of polymorphisms accrued over decades of inbreeding^38–40^; and 2.) ABX treatment can have a direct impact on eukaryotic cell function in the host^41^. To address these, we developed a colony of genetically identical C57BL/6 mice harboring CR or JAX microbiomes through vertical transfer, which we call genetically identical <u>m</u>ice colonized with <u>d</u>ivergent <u>m</u>icrobiomes at birth (IMDM; Figure 1a). 16S sequencing of fecal samples showed efficient, stable, and generally representative microbial transfer from donor mouse to IMDM host (Figure 1b-e), with *Muribaculaceae* being comparatively most abundant in IMDM-JAX mice and *Bacteroidaceae* being comparatively most abundant in IMDM-CR mice (Figure 1f, g). Consistent with our results in the commercially sourced mice (Extended figure 1), M3-9-M tumour growth was delayed, and OS extended, in female IMDM-JAX vs. -CR mice (Figure 1i, j). This difference was not observed in tumours grown in H-Y-compatible male mice (Figure 1k,l) unless the tumour cells were engineered to express a model tumour antigen (ovalbumin, OVA; Figure 1m-o). Together, these results provide direct evidence that divergent microbiomes can shape host cancer immune surveillance in ways that impact tumour control.

**Figure 1:**
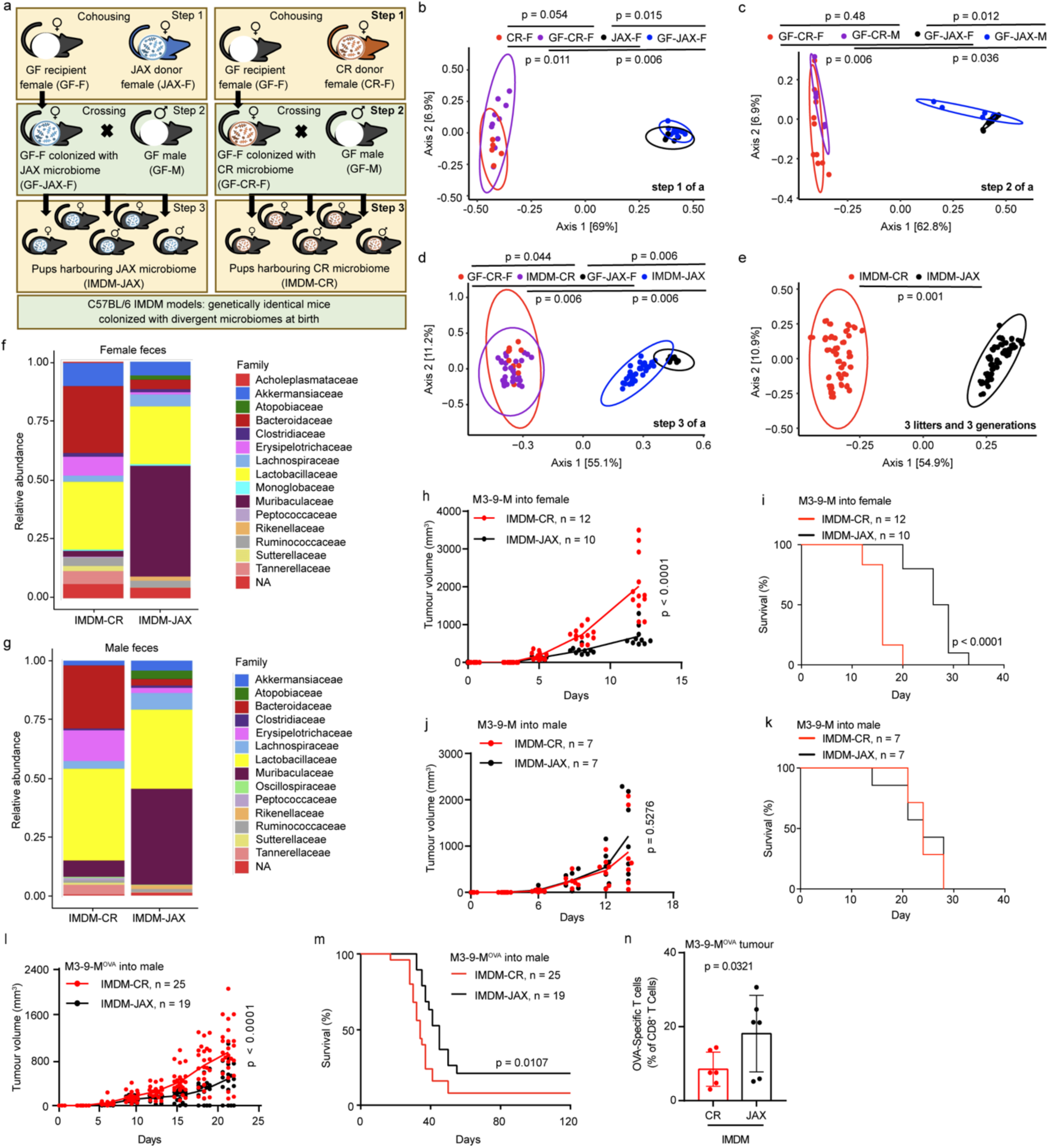
The microbiome impacts tumour immune surveillance in genetically identical hosts. a, steps followed for the production of IMDM mice; b-d, PCoA of the fecal microbiome at the end of each production step for IMDM mice (circular dot represent each mouse, PERMANOVA test, weighted UniFrac method); e, PCoA of the fecal microbiome of 6-8 week old IMDM-CR vs. -JAX mice, combining 3 litters over 3 generations (circular dot represent each mouse, PERMANOVA test, weighted UniFrac method); f-g, relative abundance of fecal bacteria in IMDM mice, identified through 16S amplicon sequencing; h-i, tumour growth kinetics and OS for M3-9-M tumours grown in IMDM-CR vs. -JAX female mice (n = 10-12/group, two combined experiments, two-way ANOVA for growth kinetics, Log-Rank test for OS); j-k, tumour growth kinetics and OS for M3-9-M tumours grown in IMDM-CR vs. -JAX male mice (n = 7/group, two combined experiments, two-way ANOVA for growth kinetics, Log-Rank test for OS). l-m, tumour growth kinetics and OS for M3-9-M^OVA^ tumours grown in IMDM-CR vs. - JAX male mice (n = 12-25/group, two combined experiments, two-way ANOVA for growth kinetics, Log-Rank test for OS); n, OVA-specific CD8^+^ T cells within M3-9-M^OVA^ tumours grown for 18 days, measured by flow cytometry (n = 6/group, two combined experiments, error bars represent standard deviation, unpaired t-test, error bars represent standard deviation).

### Divergent microbiomes create distinctive metabolomes in the TME

Commensal microbes commonly interact with host immune cells via secreted metabolites^16,32,42^. We compared fecal and tumour metabolomes of IMDM-CR and -JAX mice using untargeted, ultra-high performance liquid chromatography mass spectrometry (UHPLC-MS) and 120 metabolite standards (Extended data table 1). As predicted, IMDM-CR metabolomes were distinct from IMDM-JAX metabolomes (Figure 2a-d; Extended figure 2a-d,) and both were modifiable by ABX treatment (Extended figure 2e-h). Numerous metabolites were differentially abundant in IMDM-CR vs. -JAX mice (Figure 2e,f; Extended figure 2i,j), including L-glutamine (L-gln) and hydroxyphenyl propanoate (HPP), which were differentially abundant in fecal and tumour samples harvested from both male and female mice (Extended data table 2 and 3). The relative level of L-gln was higher in IMDM-CR mice (Figure 2g,h and Extended figure 2k,l), consistent with previous research showing that L-gln promotes tumour growth^43^. In contrast, the relative level of HPP was higher in IMDM-JAX mice (Figure 2i,j; Extended figure 2m,n), suggesting that HPP may be associated with enhanced tumour control. Importantly, ABX treatment reduced HPP levels in both feces and tumour (Figure 2k,l), suggesting that it is produced by commensal bacteria. To identify potential HPP synthesizing microbes, we performed untargeted UHPLC-MS metabolomics and 16S amplicon sequencing in parallel with predicting microbial metabolic profiles using MelonnPan^44^. We identified 77 gut bacterial amplicon sequence variants (ASVs) positively associated with HPP, including members of the Muribaculaceae family found to be correlated with enhanced antitumour immunity and highly enriched in the JAX microbiome^36^ (Extended figure 1b; Extended data table 4). Taken together, these results demonstrate that divergent microbiomes create distinct and modifiable metabolomes within host tumours, and that specific metabolites produced may influence tumour progression, potentially by impacting host antitumour immunity.

**Figure 2:**
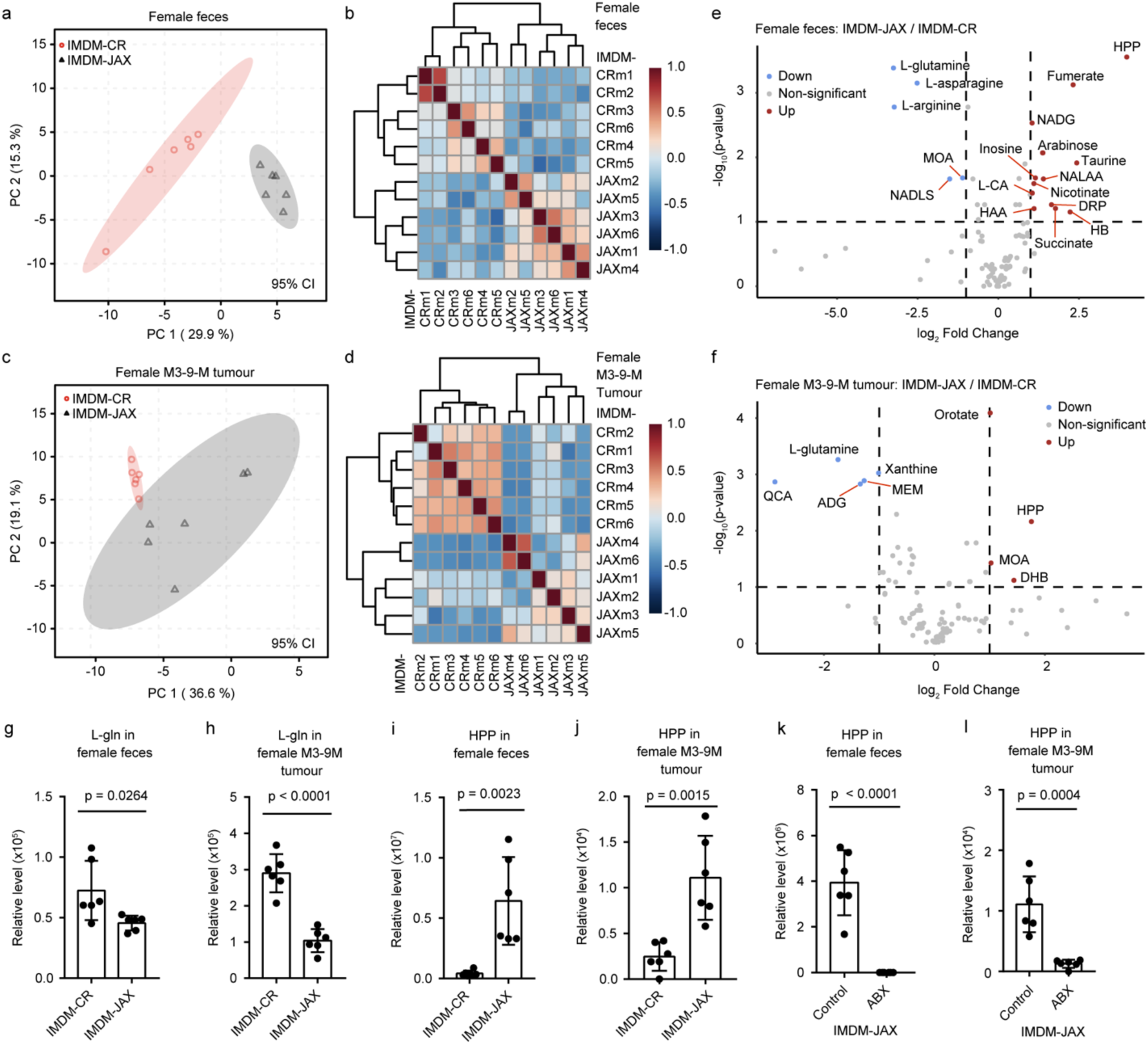
Divergent microbiomes create distinctive metabolomes in the TME. a-b, PC score and Spearman correlation heatmap of the fecal metabolome of IMDM-CR vs. –JAX female mice (samples pooled from two different experiments, ellipses drawn with 95% confidence interval); c-d, PC score and Spearman correlation heatmap of M3-9-M tumour metabolome in IMDM-CR vs. –JAX female mice (samples pooled from two different experiments, ellipses drawn with 95% confidence interval); e-f, volcano plots of fecal (e) and tumour (f) metabolites from IMDM-CR vs. –JAX female mice. Blue and red dots indicate differentially abundant metabolites, with fold change threshold (x) 2 and t-test threshold (y) 0.05 (samples pooled from two different experiments, FDR adjusted, log transformed p values); g-h, relative level of L-gln metabolite in feces and M3-9-M tumour of IMDM-CR vs. –JAX female mice (samples pooled from two different experiments, error bars represent standard deviation, unpaired t-test); i-j, relative level of HPP metabolite in feces and M3-9-M tumour of IMDM-CR vs. –JAX female mice (samples pooled from two different experiments, error bars represent standard deviation, unpaired t-test); k-l, relative level of HPP metabolite in fecal and M3-9-M tumour of IMDM-JAX female mice receiving normal vs. ABX-supplemented water (samples pooled from two different experiments, error bars represent standard deviation, unpaired t-test).

### Microbiome-derived HPP metabolites enhance anticancer immunity in mice

HPP are organic compounds belonging to the class of phenylpropanoic acids comprising a propionic acid backbone substituted with a 2-, 3-, or 4-hydroxyphenyl group at the 3-position (3,2-HPP, 3,3-HPP, and 3,4-HPP; Extended figure 3a). They are enriched in animal microbiome samples, including those from mice and humans, are metabolized by microbes from dietary precursors such as flavonoids, caffeic acid, and proanthocyanidins, and have been associated with antioxidant, anti-inflammatory, antiviral and anticancer properties^45–48^. Having found elevated HPP levels in JAX mice, which harbor a microbiome that supports anticancer immunity, we next asked whether HPP supplementation improves tumour immune surveillance and response to cancer immunotherapy. A preliminary dose-finding study in IMDM-CR mice found that 83 mg/kg of purified 3,2-HPP delivered i.p. resulted in a transient increase in [serum HPP] without causing overt toxicity or weight loss (Extended figure 3b-e). At this dose, HPP supplementation prolonged survival of IMDM-CR male mice implanted with immunogenic M3-9-M^OVA^ cells, but not those implanted with parental M3-9-M cells (Figure 3a,b). HPP supplementation similarly extended survival of IMDM-CR female mice implanted with B16.F10 melanoma cells (Figure 3c). The impact of 3,2-HPP, 3,3-HPP or 3,4-HPP supplementation was similar in all models tested. Importantly, HPP supplementation was effective against established M3-9-M and B16.F10 tumours in IMDM-CR mice, when initiated 7 days after cell implantation (Figure 3d,e). Collectively, these findings demonstrate that HPP supplementation can enhance the control of immunogenic tumours, even in hosts harbouring a microbiome that normally suppresses anticancer immunity.

**Figure 3:**
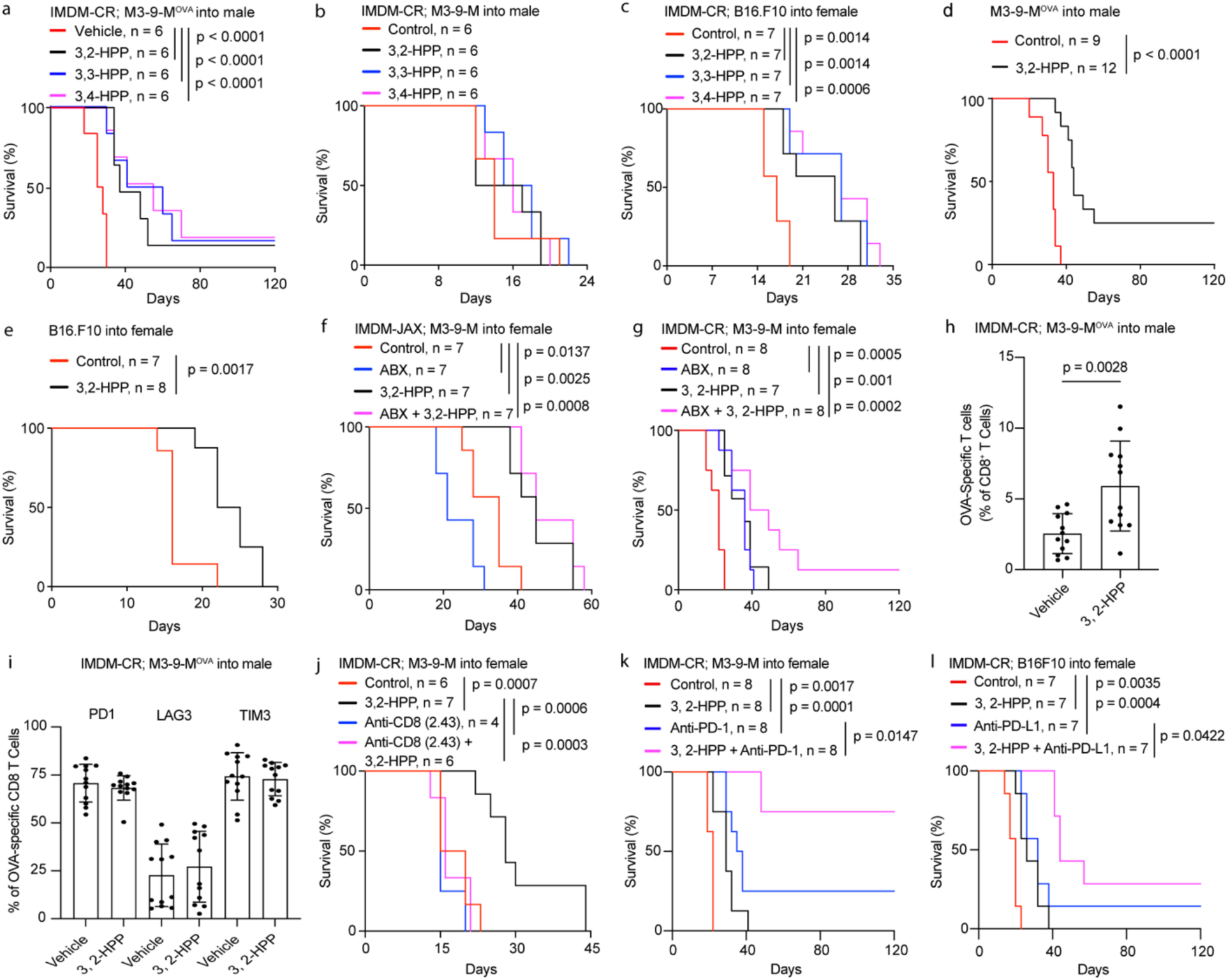
Microbiome-derived HPP metabolites enhance anticancer immunity in mice. a-c, OS curves of M3-9-M^OVA^ in male IMDM-CR, M3-9-M in male IMDM-CR, and B16.F10 in female IMDM-CR mice, respectively, receiving i.p. treatments of HPP metabolites initiated from day 1 post-cell implantation (two combined experiments, Log-Rank test); d-e, OS curves of M3-9-M^OVA^ in male IMDM-CR and B16.F10 in female IMDM-CR mice, respectively, receiving i.p. treatments of 3,2-HPP initiated from day 7 post-cell implantation (two combined experiments, Log-Rank test); f-g, effects of ABX on OS of M3-9-M in female IMDM-JAX and IMDM-CR mice, respectively, receiving i.p. treatments of 3,2-HPP from day 1 post-cell implantation (two combined experiments, Log-Rank test); h-i, flow cytometry detection of OVA-specific CD8^+^ T cell in the TME of M3-9-M^OVA^ in male IMDM-CR mice, 18 days post-tumour implantation, receiving i.p treatments of 3,2-HPP treatment from day 1 post-cell implantation, and the expression of exhaustion markers on this cell type, respectively (two combined experiments, error bars represent standard deviation, unpaired t-test); j, effects of CD8^+^ T cell depletion on OS of M3-9-M in IMDM-CR female mice, receiving i.p. treatments of 3,2-HPP from day 1 post-cell implantation (Log-Rank test, single experiment); k-l, effects of anti-PD-1 and anti-PD-L1 therapies on OS of M3-9-M and B16.F10 in female IMDM-CR mice, respectively, receiving i.p. treatments of 3,2-HPP from day 1 (two combined experiments, Log-rank test).

We next assessed the impact of HPP supplementation on mice receiving broad-spectrum ABX, which are frequently used to manage infections in cancer patients and can lead to microbiome dysbiosis^49^ and a dampened response to immunotherapy^21^. Consistent with our results in wildtype mice (Extended figure 1k,l), ABX treatment reduced M3-9-M tumour control in IMDM-JAX mice while improving tumour control in IMDM-CR mice (Figure 3f,g). HPP supplementation extended survival in all ABX-treated mice, counteracting the negative impact of ABX in IMDM-JAX mice while enhancing positive impact of ABX treatment in IMDM-CR mice. This suggests a potential benefit of HPP supplementation in a broad spectrum of cancer patients, including those undergoing ABX treatment for infection.

Our data shows that HPP-mediated antitumour activity requires tumour antigenicity, suggesting that HPP works by promoting tumour-specific CD8^+^ T cell function. Indeed, HPP supplementation increased the frequency of OVA-specific CD8^+^ T cells infiltrating M3-9-M^OVA^ tumours (Figure 3h), although it did not impact their surface expression of exhaustion markers PD-1, TIM-3, or LAG-3 (Figure 3i). Importantly, CD8^+^ T cell depletion (Extended figure 3f) abrogated the anticancer activity of HPP supplementation (Figure 3h). Together these data suggest that HPP improves anticancer immunity and tumour control by promoting antigen-specific CD8^+^ T cell accumulation in the TME without impacting their level of exhaustion. This prompted us to evaluate HPP supplementation in combination with immune checkpoint blockade (ICB) therapy, which strengthens anticancer immunity by protecting against CD8^+^ T cell exhaustion^50^. Strikingly, HPP supplementation significantly improved tumour response to PD-1/L1 blockade therapy in both M3-9-M and B16.F10 tumour models (Figure 3k,l; Extended figure 3g,h). Taken together, these data show that HPP supplementation can potentiate CD8^+^ T cell-mediated tumor immune surveillance in a manner that is complementary with ICB therapy.

### HPP metabolites potentiate innate immune signaling in tumour-infiltrating leukocytes

To identify the mechanism(s) by which HPP metabolites enhance CD8^+^ T cell-mediated antitumour immunity, we used single-cell RNA sequencing (scRNAseq) to analyze global gene expression in immune cells isolated from M3-9-M^OVA^ tumours grown in male IMDM-JAX or male IMDM-CR ± HPP. A total of 14 distinct immune cell clusters were annotated based on cell type specific gene expression markers (Figure 4a; Extended figure 4; Extended data table 6,).

**Figure 4:**
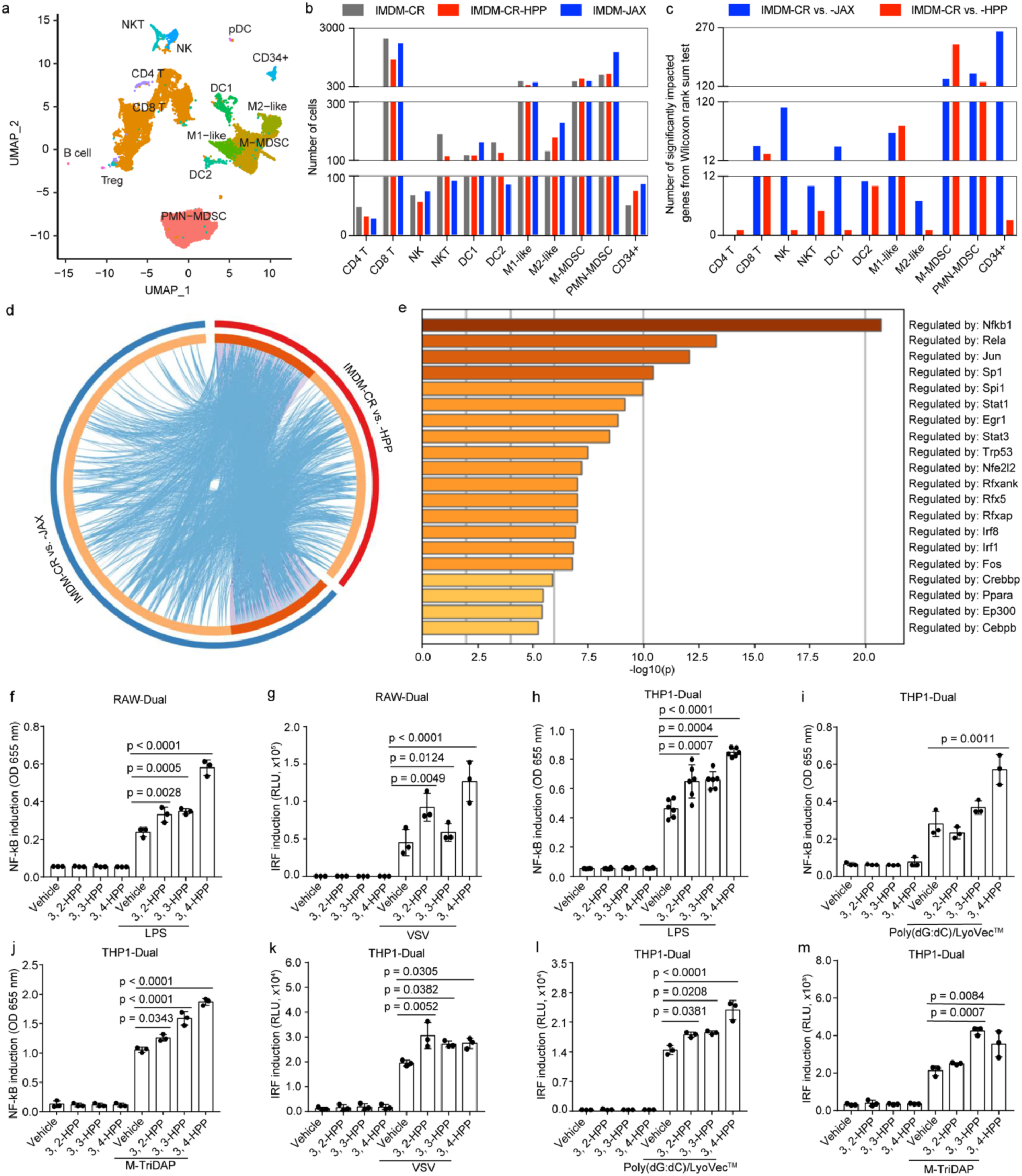
HPP metabolites potentiate innate immune signaling in tumour-infiltrating leukocytes. a, immune cell types in the TME identified through scRNAseq of CD45^+^ cells isolated from orthotopically implanted M3-9-M^OVA^ cells; b, number of immune cell types detected in M3-9-M^OVA^ tumours through scRNAseq experiment (n = 2 pooled samples per experimental groups); c, number of significantly impacted genes obtained from pairwise comparisons (Wilcoxon rank sum test); d, circos plot of all significantly impacted genes, obtained from pairwise comparisons of all cell types measured, that overlap with each other through identical or shared pathways; e, bar graph of transcriptional regulators of 3,2-HPP impacted genes; f-m, NF-κB and IRF induction in mouse RAW-Dual™ cells and human THP1-Dual™ cells with different pathogen recognition receptor agonists in the presence of HPP metabolites (HI = heat inactivated, three combined experiments, error bars represent standard error, one-way ANOVA test).

While the number of cells in each cluster was not significantly different between groups (Figure 4b), gene expression profiles were substantially altered within multiple immune cell types (Figure 4c). This was observed in pairwise comparisons between IMDM-CR vs. -JAX and also in IMDM-CR ± 3,2-HPP, in PMN-MDSC, M-MDSC, M1-like macrophages, CD8^+^ T cells, DC2, and NKT cells. Importantly, many of the genes differentially expressed between IMDM-JAX vs. IMDM-CR mice were similarly impacted by HPP supplementation of IMDM-CR mice (Figure 4d). This suggests that HPP is a major driver of communication between the JAX microbiome and tumour-infiltrating immune cells, especially in MDSC cells (Extended figure 5).

Subsequent analyses of the transcriptional regulators of HPP-impacted genes identified several pathways known to impact cancer immunity^51,52^, including NF-κB and interferon/IRF signaling (Figure 4e). Interestingly, HPP alone had no impact on NF-κB or IRF activity in a mouse monocytic cell line (RAW-Dual™; Figure 4f, g). However, when combined with stimulators of innate immune signaling, including lipopolysaccharide (LPS), vesicular stomatitis virus (VSV) and others, all three HPP metabolites enhanced NF-κB and IRF activity, with 3,4-HPP generally having the most robust impact. Importantly, a similar response was seen in a human monocytic cell line (THP1-Dual™; Figure 4h-m; Extended figure 6a-e). These results suggest that HPP metabolites potentiate innate immune signalling in myeloid cells triggered by pathogen- or danger-associated molecular patterns.

### HPP metabolites enhance GSDMD activity in myeloid cells

To learn how HPP molecules potentiate innate immune signaling, we next sought to identify its cellular binding partner(s). Using thermal proteome profiling^53^ and nonparametric analysis of the response curves (NPARC)^54^, 647 proteins were found to have heat dissociation curves altered by HPP treatment of THP1 cells (Figure 5a). When analyzed for known interactions with transcriptional regulators of HPP-modulated genes in the TME (presented in Figure 4f), GSDMD was identified as a putative HPP binder of interest (Figure 5b, c). As GSDMD is known to promote anticancer immunity in mice^57^ and associated with good outcomes in cancer patients^58^, we prioritized it for further study.

**Figure 5:**
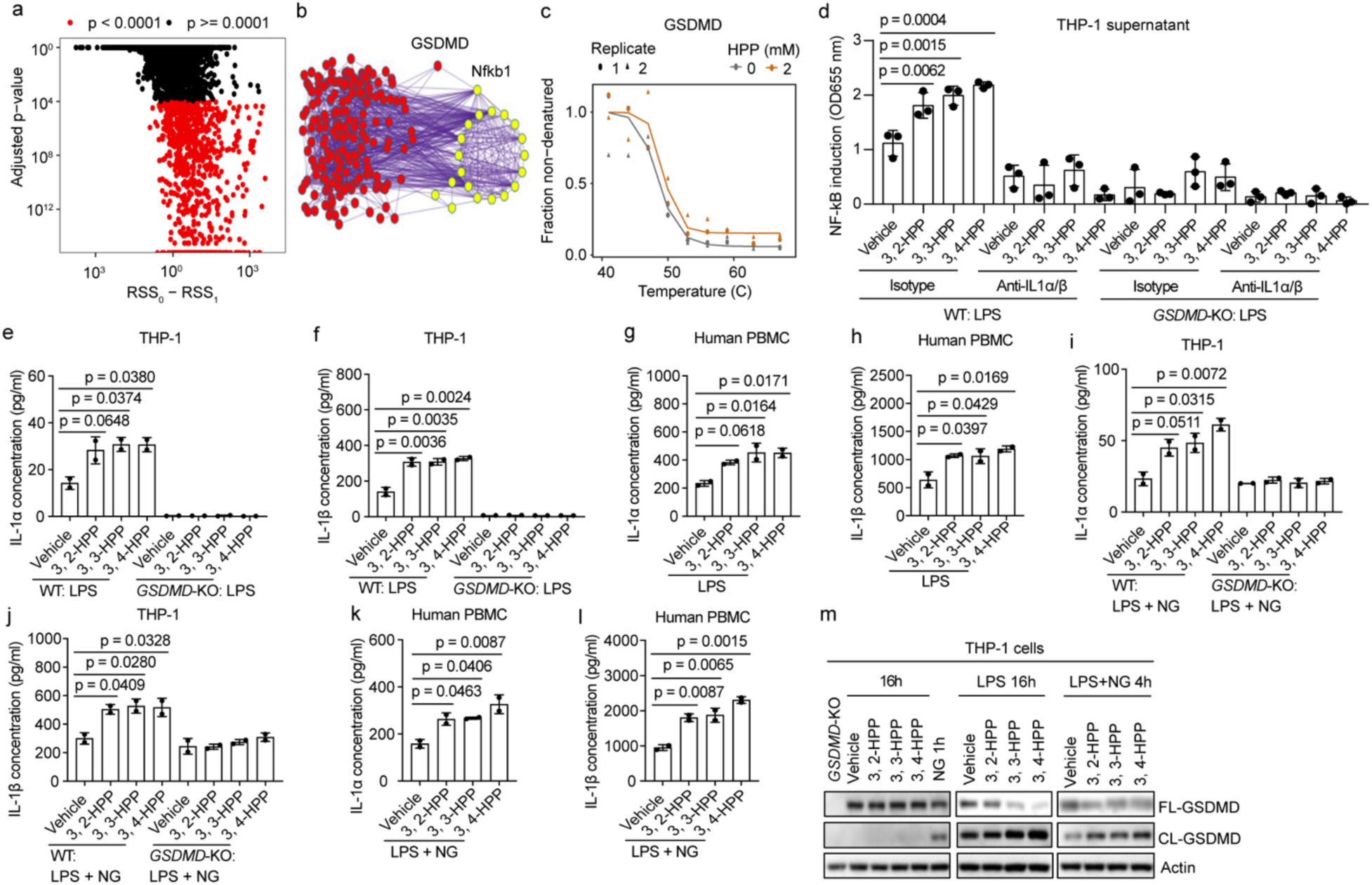
HPP metabolites enhance GSDMD activity in myeloid cells. a, representation of the proteins with denaturing curves significantly altered (red) or unaltered (black) by 3,2-HPP treatment of THP1-Dual™ cells, using thermal proteome profiling (two combined experiments, proteomic coverage: 4301, NPARC test); b, protein-protein interaction network of proteins with denaturation curves altered by 3,2-HPP treatment (red) and the transcriptional regulators of 3, 2-HPP impacted genes in M3-9-M tumours (yellow); c, impact of 3,2-HPP treatment of THP1-Dual™ cells on the denaturation curve of GSDMD (two-combined experiments, NPARC test); d, NF-κB induction in THP1-Dual™ reporter cells treated with conditioned media harvested from WT or *Gsdmd-*KO THP1 cells treated with LPS ± HPP metabolites ± IL-1α and −1β neutralizing antibody (three combined experiments, one-way ANOVA with Bonferroni multiple comparison test); e-h, [secreted IL-1α and −1β] in conditioned media harvested from THP-1 WT or *Gsdmd*-KO cells, or human PBMCs, cultured for 16 hours in LPS ± HPPs (two combined experiments, one-way ANOVA with Bonferroni multiple comparison test); i-l, [secreted IL-1α and −1β] in conditioned media harvested from THP-1 WT vs. *Gsdmd*-KO cells, or human PBMCs, cultured for 16 hours in LPS and NG ± HPPs (two combined experiments, one-way ANOVA with Bonferroni multiple comparison test); m, western blot of THP-1 cells treated with LPS ± NG ± HPPs (representative of two experiments).

GSDMD is a pore-forming protein that is highly expressed in myeloid cells and mediates the secretion of proinflammatory cytokines, such as IL-1 family members, during inflammasome activation^55,56^. Given that IL-1α and β are strong triggers of NF-κB signaling (through the IL-1 receptor, IL-1R)^59^ and that *Gsdmd*, *Nfkb1*, and other key inflammasome-related genes are enriched in tumor-infiltrating leukocytes (Extended figure 7), we hypothesized that HPP potentiates innate immune signaling by enhancing IL-1α and β secretion through GSDMD pores in myeloid cells. These molecules may in turn trigger autocrine and paracrine NF-κB and interferon signaling in myeloid and other tumour-infiltrating leukocytes, resulting in enhanced anticancer immunity.

To test this, we treated wild-type (WT) or *Gsdmd*-knockout (KO) THP1 cells with LPS±HPP in the presence or absence of neutralizing antibodies targeting IL-1α and β. As expected, media conditioned from LPS-treated WT cells (donor) induced NF-κB signaling when added to reporter cells (recipient), which was enhanced by HPP treatment of the donor cells (Figure 5d). Consistent with our hypothesis, HPP treatment enhanced IL-1α and β secretion from LPS-treated WT cells, and IL-1α and β neutralization completely blocked NF-κB induction by LPS±HPP-conditioned media (Figure 5d-f; Extended figure 8a). In contrast, HPP treatment did not increase IL-1α and β secretion from LPS-treated *Gsdmd*-KO cells or potentiate NF-κB signalling in cells treated with media conditioned from *Gsdmd*-KO cells. Interestingly, HPP treatment did not impact lactate dehydrogenase (LDH) release from LPS-treated THP-1 cells (Extended figure 8b), indicating that heightened IL-1α and β secretion is not the result of pyroptosis. Importantly, HPP treatment potentiated IL-1α and β secretion from primary human PBMC cultured with LPS (Figure 5g,h). Collectively, these data suggest that HPP metabolites strengthen innate immune signalling by potentiating IL-1α and β secretion from GSDMD pores in myeloid cells without causing cell death.

In a complementary approach, we treated THP-1 or PBMC with HPP in the presence of LPS and nigericin (NG), which together induce IL-1α and β release in a cell death dependent manner^60^. As expected, HPP enhanced IL-1α and β secretion, which in THP-1 cells was dependent on GSDMD (Figure 5i-l, Extended figure 8c,d). Unexpectedly, HPP reduced LDH release in a manner that was dependent on GSDMD (Extended figure 8e,f). Since GSDMD undergoes cleavage by inflammatory caspases upon activation^55^, we next assessed the impact of HPP on this process. While HPP molecules alone did not trigger GSDMD cleavage, their addition to LPS or LPS and NG treatment substantially enhanced GSDMD cleavage (Figure 5m). These findings indicate that HPP metabolites potentiate GSDMD activation and subsequent cytokine secretion from monocytic cells while simultaneously protecting against pyroptosis.

### GSDMD is required for HPP activity in mice and its activation is positively associated with response to ICB therapy in humans

Finally, we sought to determine if GSDMD is required for HPP to potentiate anticancer immunity in mice and how GSDMD activation is related to ICB response in human cancer patients. To begin, we treated WT or *Gsdmd*-KO mice with HPP and monitored M3-9-M^OVA^ tumour growth and survival over time. HPP did not impact tumour growth or survival in *Gsdmd*-KO mice (Figure 6a,b), indicating that HPP is dependent on host GSDMD to enhance tumour immune surveillance in mice. Consistent with our cell culture studies (Figure 5), HPP supplementation increased IL-1β levels in the interstitial fluid of M3-9-M^ova^ tumours grown in WT but not *Gsdmd*-KO mice (Figure 6c).

**Figure 6:**
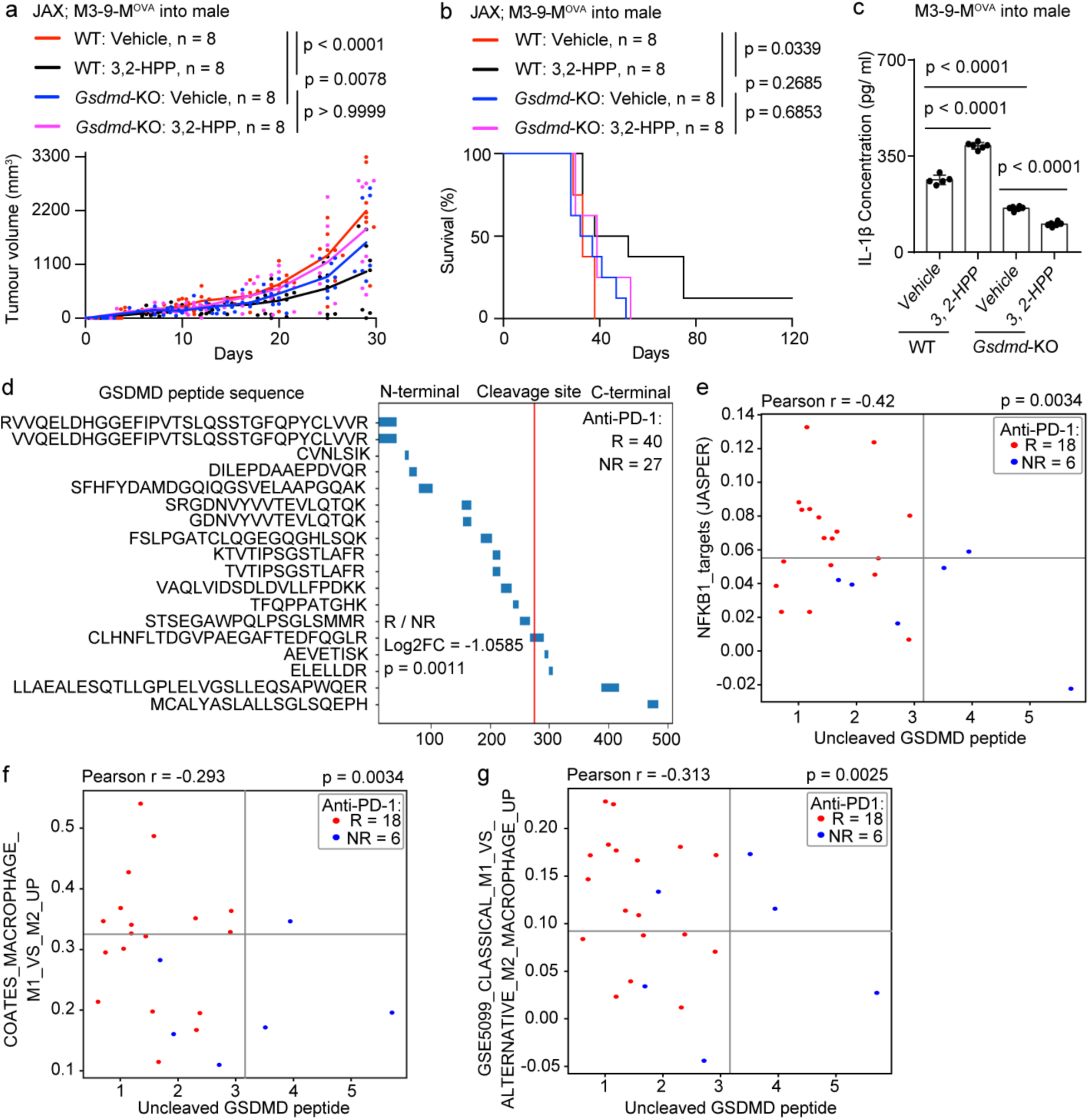
GSDMD is required for HPP activity in mice and its activation is positively associated with response to ICB therapy in humans. a-b, tumour growth kinetics and OS for orthotopic M3-9-M^OVA^ tumours in male WT or *Gsdmd*-KO mice receiving vehicle or 3,2-HPP treatments from day 1 post cell-implantation (n =8/group; two combined experiments, two-way ANOVA test for growth kinetics, Log-Rank test for OS); c, [IL-1β] in interstitial fluid harvested from 18-day old M3-9-M^OVA^ tumours grown in male WT or *Gsdmd*-KO mice receiving vehicle or 3,2-HPP treatment from day 1 post-cell implantation (two combined experiments, one-way ANOVA with Bonferroni multiple comparison test); d, GSDMD peptides quantified from a publicly available tumour proteomic database established from stage IV melanoma patients undergoing anti-PD-1 treatment (responder, R = 40 and non-responder, NR = 27); e-g, correlation uncleaved GSDMD peptide with *NFKB1* gene targets and two sets of M1- vs. M2-like macrophage gene signatures, respectively, in stage IV melanoma patients undergoing anti-PD-1 treatment (R = 18 and NR = 6, chi-square test).

Next, we explored the association of GSDMD cleavage with responses to anti-PD-1 therapy of stage IV melanoma patients, using a publicly available proteomic dataset^61^. In this dataset, the abundance of full length GSDMD peptide containing the D275 residue after the FLTD cleavage motif was lower in patients who responded to anti-PD-1 therapy (Extended data table 7; Figure 6d). Considering the pivotal role played by cleavage at the D275 residue in GSDMD activation^55,62^, these data suggests that GSDMD activity in the tumour is correlated with a favorable response to anti-PD-1 therapy. Consistent with this observation, gene set enrichment analysis (GSEA) showed a negative correlation between the abundance of full length GSDMD peptide and the expression levels of NF-κB regulated genes in this melanoma cohort (Figure 6e). Since NF-kB signaling is implicated in macrophage polarization within the TME^63,64^, we further assessed the association of GSDMD cleavage with M1 vs. M2-like macrophage polarization. Single-sample GSEA (ssGSEA) showed enrichment of M1-like macrophage-specific gene sets in anti-PD-1 responders, which inversely correlated with levels of full length GSDMD peptide (Figure 6f,g).

Collectively, our findings in cell culture, mice and humans fit a model in which HPP potentiates GSDMD activation in tumour-associated myeloid cells to improve cancer immune surveillance and response to immune checkpoint blockade therapy (Extended figure 9).

## DISCUSSION

The immune system’s ability to control cancer is complex and incompletely understood. Over the past decade, growing evidence implicates the collection of commensal microbes and their derived metabolites in modulating tumour immune surveillance and response to cancer immunotherapy^10–17^. Indeed, recent studies have demonstrated that specific microbiome-derived metabolites can boost anticancer immunity in mice^16,32,35^. Some of these metabolites act directly on anticancer T cells, for example by eliciting epigenetic or metabolic reprogramming^16,33,35^, while others work indirectly, for example by modulating the activation of tumour-infiltrating myeloid cells^31,32,65^. The latter mechanism is intriguing as myeloid cells are highly abundant in solid cancer TMEs and are at the center of immune-tumour interactions, serving as sentinels that detect cancer cells, initiate adaptive antitumour immunity in the draining lymph node, and boost anticancer T cell activity within the tumour^2,5^. However, myeloid cells within solid TMEs are commonly polarized toward immunosuppression and actively participate in immune evasion^2,66^. Owing to their abundance and plasticity in solid tumours, the molecular mechanisms governing their function are considered therapeutically actionable and are under intense investigation^2,66^.

Our study identifies a family of microbiome-derived metabolites called HPPs that modulate myeloid cell biology and promote cancer immune surveillance and response to cancer immunotherapy in mice. There are three HPP isomers, 3,2-HPP, 3,3-HPP and 3,4-HPP, each composed of a phenylpropionic acid hydroxylated at the ortho-, meta- or para-position. Produced by a variety of commensal microbes in mice and humans, HPPs are known to interact with the immune system in ways that promote anti-inflammatory, antioxidant, antiviral or anticancer functions in some contexts ^45–48^. While their molecular targets have not yet been determined, one study showed that 3,4-HPP (also called desaminotyrosine, DAT), a byproduct of dietary flavonoid metabolism within enteric bacteria, primes the amplification loop of type 1 interferon signalling and protects hosts from influenza virus pathology^45^. That mechanism prompted a recent study exploring the impact of DAT supplementation on ICB therapy in mice^34^. Similar to our findings for all three HPP isomers, those authors reported that DAT potentiates antigen-specific T cell activation and response to ICB therapy. Although a molecular mechanism was not elucidated, the impact of DAT on ICB therapy was dependent on the host type I interferon receptor, and DAT supplementation potentiated DC activation by LPS stimulation in culture^34^. The authors thus speculated that DAT promotes anticancer immunity by potentiating DC activation through amplification of type I IFN signaling.

Using unbiased thermal proteomic profiling of myeloid cells saturated with 3,2-HPP, our study identifies the first known molecular target of HPPs, the pore forming protein GSDMD. While HPPs may have several molecular targets – indeed, we identified >600 proteins with denaturation curves altered by 3,2-HPP – GSDMD was required for HPP to potentiate pattern recognition receptor (PRR) agonists-mediated cytokine release and subsequent paracrine NF-kB signal transduction in cultured myeloid cells. It was also required to improve tumour control and overall survival in mice. This suggests that host GSDMD is a primary effector of HPP-elicited antitumour activity. GSDMD is enriched in myeloid cells, where it serves as a central effector of inflammasome activation^60,67^. Its cleavage by caspase proteins activated by PRR signaling leads to multimer assembly and pore formation in the plasma membrane, through which IL-1 family and other inflammatory cytokines can shuttle. Like inflammation, GSDMD may have multiple, potentially conflicting and context-dependent roles in tumour immunity. While some reports showed that GSDMD restricts antitumour immunity during ICB therapy^67,68^, others demonstrated that its activation improves effector CD8 T cell responses to cancer^69–74^. Our data are consistent with the latter reports, as well as a recent study that identified a different small molecule GSDMD agonist called 6,7-dichloro-2-methylsulfonyl-3-N-tert-butylaminoquinoxaline (DMB) that potentiates ICB therapy in mice^57^. In that study, DMB was found to induce cleavage-independent GSDMD pore formation and cancer cell pyroptosis, an immunogenic form of cell death that elicits antitumour immunity. In contrast, we report that HPP molecules are inert on their own but potentiate GSDMD cleavage in myeloid cells that are activated by PRR signaling, which causes inflammatory cytokine secretion into the TME, without pyroptosis. Although we did not map the HPP binding site onto GSDMD, the different mechanisms used by DMB and HPP to activate GSDMD suggest distinct binding. It is intriguing to consider that two different small molecules might synergize in promoting anticancer immunity by targeting the same protein in a different way and in distinct cell types, a possibility we are currently exploring.

Our scRNAseq data show that HPP alters gene expression in numerous tumour-infiltrating immune cell types. It is likely, therefore, that HPP impacts the function of many immune cell types in solid tumours, either directly or indirectly. This is supported by studies showing that both 3,3-HPP and 3,4-HPP supplementation shifts the mouse’s enteric microbiome toward bacteria that produce immunomodulatory short chain fatty acids (SCFAs)^75^. Our scRNAseq data also identified many genes impacted by the JAX microbiome that are not affected by HPP supplementation. This suggests that there are other soluble factors from the JAX microbiome that modulate gene expression in tumour-infiltrating immune cells. Despite this complexity, we focused our mechanistic investigations on the direct effect of HPPs on myeloid cell function. Our data suggests that HPPs act directly in myeloid cells by promoting GSDMD activation and the secretion of immune-stimulating inflammatory cytokines, and that those cytokines activate both myeloid and T cells in the TME. Consistent with this mechanism, one of the cytokines impacted by HPP supplementation, IL-1β, was previously shown to enhance CD8 T-cell expansion, function, and anti-tumour immunity in mice^76,77^, although it too can have a negative impact on anticancer immunity in some contexts^78,79^. Importantly, through mining a publicly available proteomic database^61^, we found evidence of our proposed mechanism in humans, where enhanced GSDMD cleavage is associated with NF-κB target gene expression, M2 to M1-like macrophage polarization, and a favourable response to ICB therapy in patients with advanced stage melanoma.

There are several limitations to our study: 1.) the resolution of our untargeted UHPLC-MS was not high enough to identify structural differences between 3,2-, 3,3- and 3,4-HPP. As such, we do not know which isomers are enriched in the feces and tumours of JAX mice; 2.) our *16S* sequencing could not resolve commensal bacteria beyond family level. Therefore, we do not know which species or strains are enriched in JAX mice nor which produce the HPP molecules; 3.) our mechanistic experiments in mice were conducted with 3,2-HPP. While we validated the mechanism in cell culture using all three HPP isomers, as well as the functional impact of HPP treatment on anticancer immunity in mice, we did not determine whether each isomers uses the same mechanism of action *in vivo*. 4.) our mouse studies were performed in a single model of rhabdomyosarcoma, with key findings validated in a second model of melanoma. As such, we do not appreciate the full breadth of impact of our findings, although we suspect that the mechanism identified, and its therapeutic potential will be relevant to a spectrum of immunogenic solid tumours rich in immunosuppressive myeloid cells.

In conclusion, our study uncovers a mechanism whereby microbiome-derived HPP molecules regulate the strength of an immune response against cancer by potentiating GSDMD activation in tumour-infiltrating myeloid cells, therapy boosting anticancer immune function in the TME. We also demonstrate that neoadjuvant HPP supplementation is well tolerated in mice and promotes cancer immune surveillance and response to ICB therapy of solid tumours. By elucidating the molecular mechanisms underlying these observations, and validating key findings in human cells and databases, our study contributes to a deeper understanding of microbiome-host interactions in cancer and paves the way for the development of therapeutic strategies leveraging HPP, GSDMD and/or myeloid cell biology for future clinical translation. Notably, our data suggest that HPP could be utilized to improve antitumor immunity in hosts harbouring diverse microbiomes, including those receiving broad-spectrum antibiotics. This finding highlights an important potential therapeutic application of HPP since a significant percentage of cancer patients require antibiotics to fight infections, which can compromise their response to immunotherapy^49,80^.

## Methods

### Cell culture

M3-9-M cells^1^ were kindly provided by Dr. Crystal MacKall (Stanford, CA, USA). M3-9-M^OVA^ cells stably expressing chicken ovalbumin (OVA) were generated by transfection with the Super piggyBac Transposase expression vector (PB210PA; System Biosciences; Palo Alto, CA, USA) and a donor plasmid encoding OVA and a puromycin resistance gene, using Lipofectamine™ 3000 (L3000001; Thermo Fisher Scientific; Waltham, MA, USA). B16.F10 (CRL-6475) and THP-1 (TIB-202™) cells were obtained from the American Type Culture Collection (ATCC; Manassas, VA, USA). RAW-Dual™ cells (rawd-ismip), THP1-Dual™ cells (thpd-nfis), HEK-Blue^TM^ IL-1β cells (hkb-il1bv2; InvivoGen), THP1-Null2 Cells (thp-nullz) and THP1-KO-GSDMD cells (thp-kogsdmdz) were obtained from InvivoGen (San Diego, CA, USA).

M3-9-M and M3-9-M^OVA^ cells were propagated in RPMI 1640 (11875119; Life Technologies, Carlsbad, CA, USA) supplemented with 10% heat-inactivated fetal bovine serum (FBS; 1284-028; Life Technologies) and 50 μM Gibco™ 2-mercaptoethanol (21985023; Thermo Fisher Scientific). To maintain transgenic M3-9-M^OVA^ cells, 1 μg/mL Gibco™ puromycin dihydrochloride (A1113803; Thermo Fisher Scientific) was added to the culture media every other cell culture passage. B16.F10 cells were propagated in Gibco™ DMEM (11965118; Thermo Fisher Scientific) with 10% heat-inactivated FBS. RAW-Dual™ cells and HEK-Blue™ IL-1β cells were propagated in DMEM, 10% heat-inactivated FBS, 100 μg/ml Normocin™ (ant-nr-1; Invivogen), and Gibco™ Penicillin-Streptomycin (100 U/ml-100 μg/ml; 15140122; Thermo Fisher Scientific). To maintain the transgenic cell lines, 100-200μg/ml of Zeocin™ (ant-zn-05; Invivogen) was added to the growth medium every other passage. THP-1, THP1-Null2 Cells, THP1-Dual™ and THP1-KO-GSDMD cells were propagated in RPMI 1640, 2 mM L-glutamine (25030149; Thermo Fisher Scientific), 25 mM HEPES (12440061; Thermo Fisher Scientific), 10% heat-inactivated FBS, 100 μg/ml Normocin™, and Pen-Strep (100 U/ml-100 μg/ml). To maintain THP1-Dual™ cells, 10 μg/ml blasticidin (ant-bl-05; Invivogen) and 100 μg/ml Zeocin™ was added to the growth medium every other passage. To maintain THP1-KO-GSDMD, 100 μg/ml Zeocin™ was added to the growth medium every other passage. Frozen stocks were prepared for each cell line, by resuspension in growth medium with 20% FBS and 10% dimethyl sulfoxide (DMSO; D8418; Sigma-Aldrich; Oakville, ON, Canada). All cell lines were tested for mycoplasma every 8-12 weeks using Venor™ GeM Mycoplasma Detection Kit (MP0025-1KT; Sigma-Aldrich).

### Animal models

All animal experiments were performed following Canadian Council for Animal Care (CCAC) guidelines and University of Calgary (Canada) Health Sciences Animal Care Committee-approved protocols (AC16-0208 and AC20-0146) in the University of Calgary’s biohazard facility (Room B53). Age- and sex-matched mice were randomly selected for different experimental groups. C57BL/6N-*Gsdmd*^em4fcw^/J (*Gsdmd*-KO; RRID:IMSR_JAX:032410) mice were obtained from the Jackson Laboratory (Bar Harbor, ME, USA). Specific pathogen free (SPF) C57BL/6J (RRID:IMSR_JAX:000664), C57BL/6NCr (RRID:IMSR_CRL:027) and C57BL/6NTac (RRID:IMSR_TAC:B6) mice were obtained from The Jackson Laboratory (Room RB08), Charles River Laboratory Inc. (Room C62; Montreal, QC, Canada) and Taconic Biosciences (Room IBU1501C; Rensselaer, NY, USA), respectively. Germ-free (GF) C57BL/6 mice were obtained from the International Microbiome Centre (IMC; University of Calgary).

Genetically identical <u>m</u>ice colonized with <u>d</u>ivergent <u>m</u>icrobiomes at birth (IMDM) were produced through cohousing and breeding in isolated cages within a specific pathogen free (SPF) biohazard facility. To avoid cross-contamination of microbiomes between experimental groups, a controlled protocol was followed for all cage changes and mouse handling, which included glove changes and Virkon™ (055-5528; BMR; Boucherville, QC, Canada) disinfection of cage surfaces and the workstation before opening each cage. Microbiome consistency across experiments was confirmed using 16S amplicon sequencing of fecal microbial DNA samples.

### Cancer models

Tumour cells were implanted orthotopically into 6–8-week-old mice. For experiments with rhabdomyosarcoma (RMS) models, 1.5 × 10^5^ M3-9-M or M3-9-M^OVA^ cells in 50 µl of PBS were implanted orthotopically into the right gastrocnemius muscle using an insulin syringe. RMS tumour volume was calculated by subtracting the volume of the gastrocnemius muscle before tumour implantation from the volume after tumour implantation (length × width × height). For the melanoma tumour model, 5 × 10^5^ B16.F10 cells in 50 µl of PBS were implanted into the right flank using an insulin syringe. Melanoma tumour volume was determined by measuring the area of the implant site (length × width^2^ × 0.5). Tumour measurements were recorded twice weekly using digital skin calipers, from implantation day to either experimental or humane endpoints.

### Microbiome composition analysis

Fecal samples were collected directly from mice into microfuge tubes and immediately stored at −80°C for later use. Microbial DNA was extracted and purified from the fecal samples using the DNeasy PowerSoil Pro Kit (47014; Qiagen; Toronto, ON, Canada) following the manufacturer’s protocol. DNA concentration was adjusted to 25 ng/µL for all samples and subsequently sent to the Centre for Health Genomics and Informatics (CHGI, University of Calgary, Canada) for sequencing. There, a 16S V3-V4 rRNA gene amplicon library was prepared and then quantified using the Kapa qPCR Library Quantification Kit (07960140001; Roche; Laval, QC, Canada). Sequencing was conducted using the Illumina MiSeq platform with a 600 Cycle MiSeq v3 kit (MS-102-3003; Illumina Inc.; San Diego, CA, USA). The sequencing files were processed using the DADA2 R package^2^ and amplicon sequence variants (ASVs) were identified. Taxonomy was assigned to ASVs using the SILVA database (version 138)^3^ maintained by DADA2. Bacterial abundance, richness, beta diversity, and ordination analyses were conducted using the phyloseq R package^4^. Permutational analysis of variance (PERMANOVA) was performed using the “pairwise.adonis” function^5^ with 999 permutations and Bonferroni correction for multivariate analysis.

### UHPLC-MS untargeted metabolomics

Snap frozen fecal and tumour samples were stored at −80⁰C immediately after collection. On the day of metabolite extraction, 20 mg of fecal or tumour tissue or 20 µl of serum was thawed on ice. Metabolites were extracted using ice cold 50% methanol (1:50 dilution). Fecal and tumour samples were homogenized by bead beating, incubated on ice for 30 minutes, and centrifuged at 21000 ×g at 4⁰C for 10 minutes. Half of the supernatant was collected and centrifuged again for further clarification. Finally, 250 µl of supernatant from each sample was collected for mass spectrometry analysis at the Calgary Metabolomics Research Facility (CMRF; University of Calgary, Canada). Metabolomic profiling was performed using ultra-high performance liquid chromatography mass spectrometry (UHPLC-MS) on a Q Exactive™ HF Mass Spectrometer (Thermo Fisher Scientific) in negative ion full scan mode (50-750m/z) at 240,000 resolution. Metabolite separation via UHPLC was conducted using a binary solvent mixture of 20 mM ammonium formate at pH3.0 in LC-MS grade water (Solvent A) and 0.1% formic acid (%v/v) in LC-MS grade acetonitrile (Solvent B). Chromatographic separation was performed using a Syncronis™ column (Thermo Fisher Scientific) at a flow rate of 600 µL/min under the following gradient conditions: 0-2 minutes, 100% B; 2-7 minutes, 100-80% B; 7-10 minutes, 80-5% B; 10-12 minutes, 5% B; 12-13 minutes, 5-100% B; 13-15 minutes, 100% B. For all runs, the sample injection volume was 2 µL. Data processing and peak detection were performed using XCMS and MAVEN software packages^6,7^. Metabolites were identified by matching m/z signals and retention times to commercial standards. Further statistical analyses were conducted using MetaboAnalyst v.5.0^8^.

### Identifying specific metabolite-producing bacteria within the same biospecimen

Fecal samples were collected from six IMDM-CR and six IMDM-JAX female C57BL/6 mice. Each sample was divided into two parts: one for UHPLC-MS untargeted metabolomics and the other for 16S amplicon. Following initial filtering of metagenomic and metabolomic data, the MelonnPan R package^9^ was used to predict bacteria responsible for HPP metabolite production. A table of paired sequence features and microbial community metabolite abundances was used as input into the “melonnnpan.train” function, enabling an unbiased identification of specific bacteria producing metabolites within the same biospecimen.

### Antibiotics-mediated dysbiosis

A broad-spectrum antibiotic cocktail consisting of ampicillin (1 mg/mL), neomycin (1 mg/mL), vancomycin (0.5 mg/mL), and metronidazole (1 mg/mL) was added to the drinking water of the mice. To improve palatability, the concentration of metronidazole was gradually increased, reaching 1 mg/mL by day 9. Antibiotic treatment was initiated two weeks before tumour inoculation and continued throughout the experiment, with antibiotic-supplemented drinking water replaced every 3-4 days. The well-being of the mice was monitored by measuring their body weights regularly. To evaluate the effectiveness of the antibiotic treatment, real-time polymerase chain reaction (RT-PCR) was performed on fecal microbial DNA using universal oligonucleotide primers targeting conserved regions of the eubacterial 16S rRNA gene (forward primer, 5′^1320^–CCATGAAGTCGGAATCGCTAG-^1341^3′; reverse primer, 5′^1431^–ACTCCCATGGTGTGACGG-^1413^3′). Fecal microbial DNA was extracted using the DNeasy PowerSoil Pro Kit. A 20 µL PCR reaction mixture was prepared using iQ™ SYBR® Green Supermix (170882; Bio-Rad; Mississauga, ON, Canada), 300 nM of each primer, 3 ng of DNA template, and Invitrogen™ UltraPure™ DNase/RNase-Free Distilled Water (10-977-023; Thermo Fisher Scientific). RT-PCR was performed on a Bio-Rad® CFX96™ Real-Time PCR system with the following thermal cycling conditions: polymerase activation and initial denaturation at 95°C for 5 minutes, followed by 35 cycles of denaturation at 95°C for 15 seconds, and annealing and extension at 60°C for 1 minute.

### Flow cytometry

Tumours were isolated from mice and single-cell suspensions were prepared using a Mouse Tumour Dissociation Kit (130-096-730; Miltenyi Biotec; San Diego, CA, USA) following the manufacturer’s instructions. Tumours were minced into ∼2–4 mm pieces using a sterile scalpel, homogenized in a gentleMACS^TM^ C tube (130-093-237; Miltenyi Biotec; San Diego, CA, USA) containing 2 mL RPMI 1640 medium with an enzyme mix for 1 minute, and incubated at 37 °C for 40 minutes with continuous gentle shaking. After homogenizing the samples again in a second gentleMACS^TM^ C tube for 2 minutes, the homogenates were filtered through a 70 µm cell strainer and then centrifuged to obtain the single-cell suspensions. Red blood cells (RBCs) were removed using RBC lysis buffer. Leukocytes were then isolated using a Percoll gradient-based separation followed by dead cell staining using Zombie Aqua™ Fixable Viability Kit (423102; BioLegend; San Diego, CA, USA). For T lymphocyte surface marker staining, the following fluorophore-conjugated antibodies were used: PE/Cy7 anti-mouse CD8a (clone 53–6.7; 100722; BioLegend), Brilliant Violet 421™ anti-mouse CD3e (clone 145-2C11; 100335; BioLegend), APC/Cy7 anti-mouse CD45 (clone 30-F11; 103116; BioLegend), Alexa Fluor® 647 anti-mouse CD279 (PD-1; clone 29F.1A12; 135230; BioLegend), PE anti-mouse CD366 (Tim-3; clone RMT3-23; 119704; BioLegend) and PerCP/Cyanine5.5 anti-mouse CD223 (LAG3; clone C9B7W; 125212; BioLegend). For OVA-specific CD8^+^ T cell detection, AF647-labelled H-2K(b) chicken ovalbumin 257-264 SIINFEKL tetramers were used. Following staining, cells were washed with FACS buffer and analyzed using an Attune NxT Flow Cytometer (Thermo Fisher Scientific).

### Mouse treatment regimens

3,2-HPP (393533; Sigma-Aldrich), 3,3-HPP (91779; Sigma-Aldrich) and 3,4-HPP (H52406; Sigma-Aldrich) were dissolved in UltraPure™ DNase/RNase-Free Distilled Water (Invitrogen™) with 1.0N NaOH (S2770; Sigma-Aldrich) to prepare stock solutions. The pH of the stock solutions was adjusted 7.4–7.6, and endotoxin levels were assessed using Pierce™ LAL Chromogenic Endotoxin Quantitation Kit (88282; Thermo Fisher Scientific; Extended data table 5). The maximum tolerable dose and pharmacokinetics of the metabolites were first established before conducting animal experiments. To test the metabolite effects on antitumour immunity, 100 µl of the metabolite (83 mg/ kg body weight) was administered i.p. every 48 hours for seven doses. For ICB experiments, mice were treated with *InVivo*MAb anti-mouse PD-1 (clone RMP1-14, BE0146; Bio X cell; Lebanon, NH, USA) and *InVivo*MAb anti-mouse PD-L1 (clone 10F.9G2, BE0101; Bio X cell) antibodies. Each antibody was administered at 250 µg per dose (i.p.) on day 10, 13 and 16 post-tumour implantation. To deplete CD8^+^ T cells, mice were treated with *InVivo*MAb anti-mouse CD8a monoclonal antibody (clone 2.43; BE0061; Bio X cell). 200 μg of anti-mouse CD8a antibody was injected i.p. per mouse on days −7, 0, 7, 14, 21, 28 and 35, where day 0 represents the day of tumour implantation. For flow cytometry, a different clone of anti-mouse CD8a antibody conjugated with PE/Cyanine7 (clone 53–6.7; 100722; BioLegend) was used for staining.

### NF-κB and IRF activity reporter assay

NF-κB and IRF induction assays were performed using mouse RAW-Dual™ and human THP1-Dual™ reporter cell lines. Cells were resuspended in freshly prepared test media at a concentration of 1.0 × 10^6^ cells/mL for RAW-Dual™ and 5.0 × 10^5^ cells/mL for THP1-Dual™. A total of 180 µL of cell suspension was seeded per well in a 96-well plate and PRR agonist concentrations were optimized for NF-κB and IRF pathway activation. GFP-expressing vesicular stomatitis virus delta M51 (VSV)^10^ was obtained from Dr. David Stojdl (CHEO Research Institute, Ottawa, ON, Canada). For stimulating RAW-Dual™ cells, LPS (100 ng/mL; tlrl-peklps; InvivoGen) or VSV (1 MOI) were used. For stimulating THP1-Dual™ cells, LPS (10 ng/mL; InvivoGen), TNF-α (0.05 ng/mL; rcyc-htnfa; InvivoGen), Poly (dA:dT)/lyovec™ (2.5 µg/mL; tlrl-patc; InvivoGen), Poly(dG:dC)/lyovec™ (2.5 µg/mL; tlrl-pgcc; InvivoGen), M-TriDAP (5 µg/mL; tlrl-mtd; InvivoGen), 2’3’-cGAMP (10 µg/mL; tlrl-nacga23-02; InvivoGen), NG (5 µg/mL; tlrl-nig; InvivoGen) or VSV (1 MOI) were used. To assess the effect of the metabolites on these pathways, 10 µL of the optimized receptor agonist and 10 µL of the 2 mM metabolite solution or vehicle were added to each well, making a final volume of 200 µL. To neutralize IL-1 receptor signalling, the following antibodies were used: anti-human IL-1α (1 µg/mL; clone 7D4; mabg-hil1a-3; InvivoGen), anti-human IL-1β (1 µg/mL; clone 4H5; mabg-hil1b-3; InvivoGen) and IgG1 isotype control (1 µg/mL; clone T8E5; mabg1-ctrlm; InvivoGen). NF-κB activity was assessed by measuring secreted embryonic alkaline phosphatase (SEAP) levels using QUANTI-Blue™ solution (rep-qbs2; InvivoGen). A 20 µL aliquot of cell culture supernatant was mixed with 180 µL of QUANTI-Blue™ and SEAP levels were determined by monitoring the colorimetric change at 650 nm using a SpectraMax i3 Microplate Reader (Molecular Devices, San Jose, CA, USA). To assess IRF induction, secreted lucia luciferase levels were measured in the culture medium using QUANTI-Luc™ (rep-qlc2; InvivoGen). A 20 µL aliquot of cell culture supernatant was mixed with 50 µL of QUANTI-Luc™ solution and luminescence was recorded immediately using the SpectraMax i3.

### Cell viability

Cell viability was assessed using the Cytotoxicity Detection Kit^PLUS^ (LDH; 4744926001; Roche) release assay, in which 46 μl of reaction mixture was added to 46 μl of cell culture supernatant in a 96-well plate. The plate was incubated in the dark for 20 minutes, and absorbance readings were recorded using the SpectraMax i3. Positive controls consisted of supernatant from cells treated with lysis solution, while negative controls were culture medium without cells.

### Cytokine assays

Cells were seeded in a 96-well plate at a density of 0.5-1.0 × 10^6^ cells/well and treated with metabolites, either alone or in combination with PRR agonists. Culture supernatants were collected at specific timepoints to measure cytokine levels. To assess cytokine levels in tumour interstitial fluid, samples were prepared as previously described^11^. IL-1 levels were determined by incubating HEK-Blue™ IL-1β reporter cells with collected culture supernatants and measuring SEAP levels in the culture medium using QUANTI-Blue™ solution. Multiplexed quantification of cytokines, chemokines, and growth factors was performed by Eve Technologies Corp. (Calgary, AB, Canada) using the Luminex™ 200 system (Luminex, Austin, TX, USA). Additionally, IL-1 alpha ELISA kit (BMS243-2; Thermo Fisher Scientific) and IL-1 beta ELISA kit (KHC0011; Thermo Fisher Scientific) were used for measuring secreted IL-1α and IL-1β levels.

### Single cell RNA sequencing (scRNAseq)

M3-9-M^OVA^ RMS cells were orthotopically inoculated into male IMDM mice and i.p. treatment of 3,2-HPP was initiated one day post-tumour implantation. The final treatment dose was administered six hours before tumour extraction. After 14 days, tumours were extracted and minced into small pieces (∼2-4 mm in diameter) using a sterile scalpel. Single-cell suspensions were prepared using Mouse Tumour Dissociation Kit (Miltenyi Biotec). Viable cells were enriched using Dead Cell Removal Kit (130-090-101; Miltenyi Biotec), and CD45^+^ tumour-infiltrating immune cells were isolated using Mouse CD45 (TIL) Microbeads (130-110-618; Miltenyi Biotec). The purity of the isolated CD45^+^ cell population was verified using flow cytometry, achieving an average purity of >95%. Cell viability was confirmed via microscopy. To enrich cell numbers for single-cell RNA (scRNA) library construction, two samples from each experimental groups were pooled. scRNA libraries were constructed using the Chromium Single Cell 3ʹ GEM, Library & Gel Bead Kit v3 (PN-1000092; 10x Genomics, Pleasanton, CA, USA) and the Chromium Controller platform (10x Genomics), with a target of 5,000 cells per library. Libraries were sequenced using the NovaSeq™ 6000 platform (Illumina) at the CHGI. A single sequencing cartridge from the NovaSeq 6000 S4 Reagent Kit v1.5 (300 cycles) (20028312; Illumina) was used, targeting 100,000 reads per cell.

### scRNAseq data analysis

Raw sequencing files were processed using Cell Ranger 6.0.2 (10x Genomics). Raw base call (BCL) files were demultiplexed into FASTQ files using the “cellranger mkfastq” command with the following sample indices: IMDM-CR = SI-GA-A3, IMDM-CR-HPP = SI-GA-B3 and IMDM-JAX = SI-GA-C3. The “cellranger count” pipeline was used for sequence alignment, filtering, barcode counting, unique molecular identifier (UMI) counting, and mapping to the 10x Genomics pre-built mouse reference genome (mm10, GENCODE vM23/Ensembl 98, 7 July 2020). To ensure equal sequencing depth across samples, the “cellranger aggr” pipeline was used for library aggregation, followed by analysis in Loupe Browser 5.1.0 (10x Genomics). Additionally, Seurat R package^12–14^ was used for scRNAseq analysis, yielding results consistent with loupe Browser outputs. The data were filtered to remove cells with low or high feature counts (<200 or >7500) and those with high mitochondrial gene frequency (>5%). After filtering, all samples were integrated into a single Seurat object, normalized, and scaled before performing both linear and non-linear dimensionality reduction. Cell clustering and annotation were conducted using cell type-specific expression markers (Extended data table 6). Differential gene expression analysis was performed using the Wilcoxon rank-sum test. To further investigate the biological significance of differentially expressed genes, Metascape^15^ and Cytoscape^16^ were used for pathway identification and protein network visualization.

### Thermal proteome profiling using intact cells

Thermal proteome profiling of intact THP1-Dual™ cells was performed by treating 24 mL of cells at a density of 1.5 × 10^6^ cells/mL with 2 mM metabolite or vehicle control for 16 hours at 37⁰C with 5% CO_2_. Cells were centrifuged at 340 ×g for 5 minutes at 4°C, resuspended in 20 mL of ice-cold PBS, and centrifuged again. The cell pellet was resuspended in 1200 µL of PBS, divided into 10 aliquots of 100 µL in 0.2 mL PCR tubes, and centrifuged at 325 ×g for 2 minutes at 4°C. After removing 80 µL of the PBS supernatant, tubes were gently tapped to resuspend the cells. Cells were subjected to heat treatment in a PCR machine for 3 minutes at temperatures ranging from 37°C to 67°C in parallel for both the metabolite- and vehicle-treated samples. Cells were then incubated at room temperature for 3 minutes. Next, cells were snap-frozen in liquid nitrogen for 1 minute, thawed briefly in a 25°C water bath, placed on ice, and resuspended by pipetting. This freeze-thaw step was repeated once, followed by the addition of 30 µL of PBS. Samples were centrifuged at 100,000 ×g for 20 minutes at 4°C, and the supernatant was transferred into a fresh tube for further processing. Protein concentration was measured for the 37°C sample, and lysates were reduced using 10 mM DTT at room temperature for 30 minutes before being alkylated with 50 mM CAA for 30 minutes in the dark. The volume of lysate corresponding to 200 µg of protein at the 37°C was determined and used as a reference to normalize protein extraction volumes across all temperature points.

### Protein clean-up and peptide digestion

Proteins were cleaned, and peptides were digested following previously described methods^17,18^. Briefly, the lysate volume corresponding to 200 µg of protein at the 37°C temperature point was adjusted to 190 µL using 50 mM HEPES buffer (J16924.K2; Thermo Fisher Scientific) for each sample. SP3 beads (Sera-Mag SpeedBeads; 41552105050250; Avantor Sciences, Radnor, PA, USA; and 65152105050250; Fisher Scientific; Hampton, NH, USA) were washed twice with 180 µL of H_2_O, resuspended in 10 µL of H_2_O, and added to the sample along with 200 µL of 100% ethanol. The sample was then incubated on a thermomixer at room temperature for 10 minutes at 1000 rpm. Following incubation, the samples were washed four times with 180 µL of 80% ethanol and the beads were resuspended in 200 µL of 50 mM HEPES buffer containing 4 µg of Trypsin/Lys-C Mix (V5071; Promega; Madison, WI, USA). Peptide digestion was carried out overnight at 37°C on a shaking platform at 600 rpm. The next day, digested peptides were transferred to a fresh Eppendorf tube, and peptide concentration was measured at the 37°C temperature point.

### Tandem mass tag (TMT) labelling and peptide fractionation

TMT labelling was performed by dissolving 0.8 mg of TMT reagents (90110; Thermo Fisher Scientific) in 80 µl of 100% Acetonitrile (ACN; 34851; Sigma-Aldrich). For each sample, an equivalent volume containing 100 µg of peptides from the 37⁰C temperature point was transferred into a fresh Eppendorf tube. TMT label (10 µL) was added to each sample, followed by incubation at room temperature for 30 minutes. This labelling step was repeated with an additional 10 µl of TMT reagent to ensure complete labelling. The reaction was quenched using 10 µl of 1 M glycine (67419; Sigma-Aldrich). All 10 labeled samples were pooled together and dried down using a SpeedVac concentrator (SPD130DLX-115; Thermo Fisher Scientific). 50% of the pooled sample was desalted using Pierce™ Peptide Desalting Spin Columns (89852; Thermo Fisher Scientific) and resuspended in high-pH mobile phase A (20 mM ammonium formate; 70221; Sigma-Aldrich). Peptide fractionation was carried out using High pH C18 Reverse Phase HPLC with mobile phase B (100% ACN; Sigma-Aldrich) under a 54-minute gradient (5%-80% B) at a flow rate of 0.2 mL/min. The 54 collected fractions were concatenated into 18 final fractions, which were then dried down. Peptides were resuspended in 20 µl of 0.1% formic acid (159013; Sigma-Aldrich) for mass spectrometry analysis.

### Mass spectrometric analysis of fractionated peptides

1 μg of peptides from each fraction was loaded onto an Orbitrap Fusion Eclipse instrument coupled to an EASY-nLC 1200 pump (Thermo Scientific). Peptides were accumulated on an Acclaim PepMap 100 C18 peptide trap column (75 um inner diameter, 2 cm length, Thermo Scientific) and then separated on an EASY-Spray HPLC Column (50cm length, 2 μm particle size and 75 μm inner diameter, Thermo Scientific). The flow rate and column temperature were kept at 250 nl/min and 40 °C, respectively. A 65 minutes gradient of 80% acetonitrile in 0.1% formic acid consisting of three linear gradient steps – from 5 to 10% B over 5min, 10 to 35% over 50 min and 35 to 95% over 10 min was used to separate the peptides. The mass spectrometer was operated in Cycle Time mode where each MS^1^ orbitrap scan (120K resolution, 375-1600 m/z scan range, 50% normalized AGC target, 30 msec maximum injection time) was followed by as many ion trap MS^2^ scans as possible as possible in a 3 sec time window, for intensities between 5000 and 1E+20 (precursor fit error maximum 50%). Ions with a charge state of 2-6 only were selected for fragmentation (32% HCD fragmentation energy, 0.7 Th isolation window, turbo scan rate, 20 msec maximum injection time, scan range 200-1400, first mass scan range mode starting at 110 m/z). Dynamic exclusion was set to 60 sec with a 10 ppm tolerance. For MS^3^, the Real-Time Search filter was included to only allow identifiable PSMs for fragmentation by the synchronous precursor selection mode (SPS). The SwissProt human proteome database (downloaded 2022-03-02) was used to perform the real-time searches. The maximum allowed number of trypsin missed cleavages was set to 1. Methionine oxidation (+15.995 Da) was set as a dynamic modification and a maximum two modifications were allowed per peptide.

Carbamidomethylation (+57.021 Da) of cysteines and TMT 6plex (+229.163 Da) on any N-terminus or lysine were set as static modifications. Maximum mass error was set to 10 ppm with fit quality parameters Xcorr of 1.4 and οCn of 0.1. Searches exceeding 35 msec were aborted. The SPS mode was then used to select the 10 most intense fragments from the MS^2^ using isobaric tag exclusion filtering with TMTpro as the reagent type (precursor exclusion with 25 ppm tolerance). Ions were fragmented simultaneously and analyzed as follows: MS^3^ isolation window 2 Th, HCD fragmentation energy 65%, resolution 50K, scan range 100-500, normalized AGC target 200% and maximum injection time 105 msec.

The data were processed using Proteome Discoverer 2.5 (Thermo Scientific). Results were searched with the Sequest algorithm against the SwissProt human proteome database (downloaded 2022-03-02) combined with the common contaminants included in MaxQuant 2.1.0.0. The combined database was concatenated with its reversed sequences for FDR calculation and search allowed for 2 missed cleavages and peptide length 6-144. Methionine oxidation (+15.995 Da), N-terminal acetylation (+42.011 Da) and N-terminal methionine loss (−131.040 Da) were set as dynamic modifications. Carbamidomethylation (+57.021 Da) of cysteines and TMT 10plex (+229.163 Da) on any N-terminus or lysine were set as static modifications. The maximum allowed mass error was 10 ppm for precursors and 0.6 Da for fragments. Identified peptides and proteins were filtered for maximum 1% FDR (Percolator algorithm). Proteins were quantified by summing reporter ion abundances across all matching PSMs belonging to unique peptides. PSMs were filtered for average reporter S/N (5 or greater). Proteins with a quantification value in every sample and possessing 2 or more unique peptides were considered quantified, normalized to the corresponding 37°C sample and their thermal profiles were evaluated by using NPARC R package^19^.

### Western blotting

Cells were collected, and total protein lysates were prepared using lysis buffer (50 mM Tris-HCl, pH 8.0; 150 mM NaCl; 1% Triton X-100; 1% SDS), and lysates were boiled at 100°C for 5-10 minutes. Protein quantification was performed using the Bio-Rad DC™ Protein Assay Kit (5000112; Bio-Rad). Proteins were separated using 12% SDS-PAGE gels and transferred to a PVDF membrane using the Trans-Blot Turbo™ Transfer System (1704150; Bio-Rad) at 25 V/2.5 A for 30 min. Membranes were blocked with Tris-buffered-saline with Tween-20 (TBST) containing 5% skim milk for 30 minutes at room temperature, followed by overnight incubation at 4°C with primary antibodies. The following antibodies were used: rabbit monoclonal recombinant anti-GSDMD antibody (ab210070; Abcam; Cambridge, UK) for full-length GSDMD (FL-GSDMD), anti-cleaved N-terminal GSDMD antibody (ab215203; Abcam) for cleaved GSDMD (CL-GSDMD) and mouse monoclonal anti-α-actin-1 (ACTA) antibody (MAB1501, Sigma-Aldrich) as a loading control. The next day, membranes were washed three times with TBST and then incubated for 1 hour at room temperature with secondary antibodies: goat anti-rabbit horseradish peroxidase (HRP)-conjugated IgG (1706515, Bio-Rad) or goat anti-mouse HRP-conjugated IgG (1706516, Bio-Rad). Secondary antibodies were diluted 1:5000 in TBST containing 5% (w/v) skim milk, followed by three washes with TBST. Protein detection was performed using the Clarity™ Western ECL Substrate (1705060; Bio-Rad) and signals were captured on a Chemidoc-It™ Imager (UVP, Upland, CA, USA).

### Experiment with human PBMCs

PBMCs were isolated from healthy adult volunteers using venipuncture to collect blood into BD Vacutainer® Venous Blood Collection Tubes (CABD366643L; BD; Mississauga, Ontario). The collected blood was diluted 1:1 with sterile PBS and layered onto Ficoll® Paque Plus (GE17-1440-02; Sigma-Aldrich) in a centrifuge tube. After centrifugation, the PBMC layer was carefully removed, washed twice in PBS, and resuspended for counting and viability assessment. PBMC treatments were performed by stimulating cells with LPS (1 ng/mL, InvivoGen) followed 3 hours later by NG (1 µg/mL, InvivoGen) in the presence or absence of HPP molecules at 37⁰C with 5% CO_2_. After 16 hours, cell supernatants were collected for cytokines and LDH measurements.

### Clinical proteomic data analysis

GSDMD peptide quantification was performed using a publicly available proteomic dataset derived from clinical samples of advanced-stage melanoma patients undergoing anti-PD-1 treatment. Peptides from both the N- and C-terminal domains were quantified, along with an uncleaved peptide spanning the GSDMD cleavage site.

Differential peptide abundance was assessed using Welch’s t-test between responders and non-responders. To assess whether targets of the NFKB1 transcription factor were upregulated in responders, we obtained a NFKB1 target gene set from JASPAR transcription factor database and calculated ssGSEA normalized enrichment scores for all samples. We then correlated the gene set scores with GSDMD uncleaved peptide abundance, and defined quadrants using the midpoints. Significance was assessed using the chi-squared test. Additionally, to assess M1 vs. M2 polarization, two gene sets were used from MSigDB: COATES_MACROPHAGE_M1_VS_M2_UP and GSE5099_CLASSICAL_M1_VS_ALTERNATIVE_M2_MACROPHAGE_UP.

### Sample size determination and statistical analysis

Where possible, sample size was predetermined based on pilot studies to estimate effect size. ClinCalc.com’s power analysis tool (https://clincalc.com/stats/samplesize.aspx) was used to estimate the required sample sizes for animal experiments. Random allocation of mice to experimental groups was ensured, and they were housed in multiple cages to eliminate cage effects. Additionally, prior experimental experience and expert consultations were used to determine sample sizes for specific techniques, including *in vitro* assays, scRNAseq, metabolomics, and thermal proteome profiling experiments. All data points were included in the analyses. Statistical evaluations of tumour growth kinetics, animal OS, and *in vitro* experiments were performed using Prism 10 (GraphPad Software, San Diego, CA, USA). Statistical analyses for thermal proteome profiling, scRNAseq, and 16S amplicon sequencing data were conducted using R software packages. To determine data distribution, either Shapiro-Wilk normality test or D’Agostino-Pearson omnibus normality test was performed. If data followed Gaussian (normal) distribution, parametric statistical tests were applied, otherwise, non-parametric tests were used. The confidence interval was set at 95%, and the specific statistical tests used, along with corresponding p-values, were reported in the respective figure or table legends.

## Acknowledgements

We thank Crystal MacKall (Stanford, CA, USA) and David Stojdl (CHEO Research Institute, Ottawa, ON, Canada) for providing M3-9-M cells and GFP-expressing vesicular stomatitis virus delta M51 (VSV), respectively. We are grateful to the NIH Tetramer Core Facility for profiding AF647-labelled H-2K(b) chicken ova 257-264 SIINFEKL tetramers.

## Author contributions

SS and DJM conceived and led the study. SS, JR, KKE, FZ and DJM wrote the manuscript, with input from all authors. SS, DK, TW, and SB conducted in vitro and in vivo preclinical studies, supervised by DJM. PD and RF performed thermal proteome profiling experiments, supervised by DS. TV analyzed the human clinical data, supervised by SM. KIE prepared libraries for single cell sequencing, supervised by MG. MD performed metabolomics, supervised by IL. KM provided germ-free mice and intellectual input. DD provided intellectual input. SS analyzed the thermal proteome profiling, scRNAseq, 16S amplicon sequencing and metabolomics data. DJM oversaw the project.

## Competing interest declaration

The authors listed in the USPTO 63/453,947 declare a potential competing interest as they have filed a patent application related to the technology described in this article. The authors declare that this patent application does not alter their adherence to the journal’s policies on data sharing and materials availability.

## Data and code availability

The data supporting the findings of this study are available within the article, its Supplementary Information files, or the referenced public repositories. The Gene Expression Omnibus (GEO) dataset for the scRNAseq experiment is available under accession code GSE232748. The Sequence Read Archive (SRA) dataset for the microbiome characterization experiment is available under accession code PRJNA974566. The Proteomic dataset for the Thermal Proteome Profiling experiment is available in the PRIDE repository under accession code PXD045790.

## Funding acknowledgements

SS was supported by the Eyes High Doctoral Recruitment Scholarship, Mitacs-Accelerate Graduate Research Internship Program, and ACHRI Graduate Scholarship. DJM was supported by the Cancer Research Society (grant number 24105), the Canadian Cancer Society (CCS) [grant number 2020–707161], and generous community donors through the Alberta Children’s Hospital Foundation (ACHF), Believe in the Gold Foundation, Alberta Cancer Foundation (ACF) and the University of Calgary.

## Extended data figure/table

**Extended figure 1:**
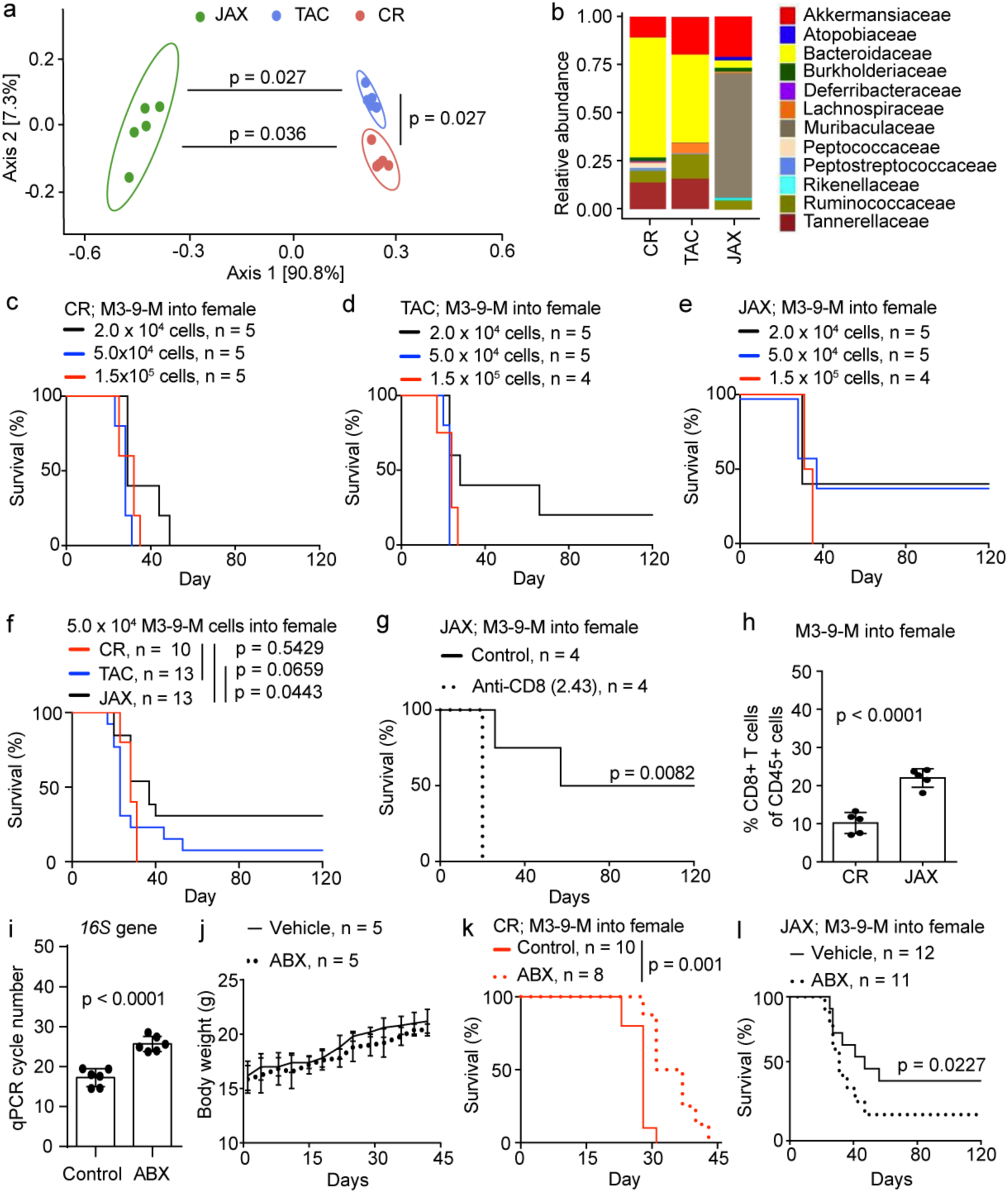
The host microbiome is associated with its capacity for tumour immune surveillance. a-b, PCoA and relative abundance of microbiota at family level in fecal microbiome samples of CR, TAC or JAX C57BL/6 female mice before tumour implantation (n = 5/ group, two combined experiments, weighted UniFrac distance, pairwise PERMANOVA test); c-e, OS of female C57BL/6 mice sourced from CR, TAC or JAX orthotopically implanted with M3-9-M tumour cells into the gastrocnemius muscle (n = 4 - 5/ group, single experiment); f, OS of CR, TAC and JAX C57BL/6 female mice orthotopically implanted with 5.0 × 10^4^ M3-9-M tumour cells (n = 10 - 13/group, two combined experiments, Log-Rank test); g, OS of JAX C57BL/6 mice orthotopically implanted with 5.0 × 10^4^ M3-9-M tumour cells and treated with or without anti-CD8 (clone 2.43) neutralizing antibody (n = 4/group, single experiment); h, CD8^+^ T cell abundance by flow cytometry in M3-9-M tumours 18 days after orthotopic implantation of 5.0 × 10^4^ cells (n = 5/ group, single experiment, error bars represent standard deviation, unpaired t-test); i, qPCR cycle number required to reach critical threshold of 16S gene expression for fecal microbial DNA in mice receiving drinking water supplemented with or without ABX for two weeks (n = 6/group, two combined experiments, error bars represent standard deviation, unpaired t-test); j, body weight of mice receiving drinking water supplemented with or without ABX (n = 5/ group, two combined experiments); k, OS of CR C57BL/6 female mice orthotopically implanted with 5.0 × 10^4^ M3-9-M tumour cells and receiving normal or ABX-supplemented drinking water (n = 8 - 12/group, two combined experiments, Log-Rank test); l, OS of JAX C57BL/6 female mice orthotopically implanted with 5.0 × 10^4^ M3-9-M tumour cells and receiving normal or ABX-supplemented drinking water (n = 11-12/group, two combined experiments, Log-Rank test).

**Extended figure 2:**
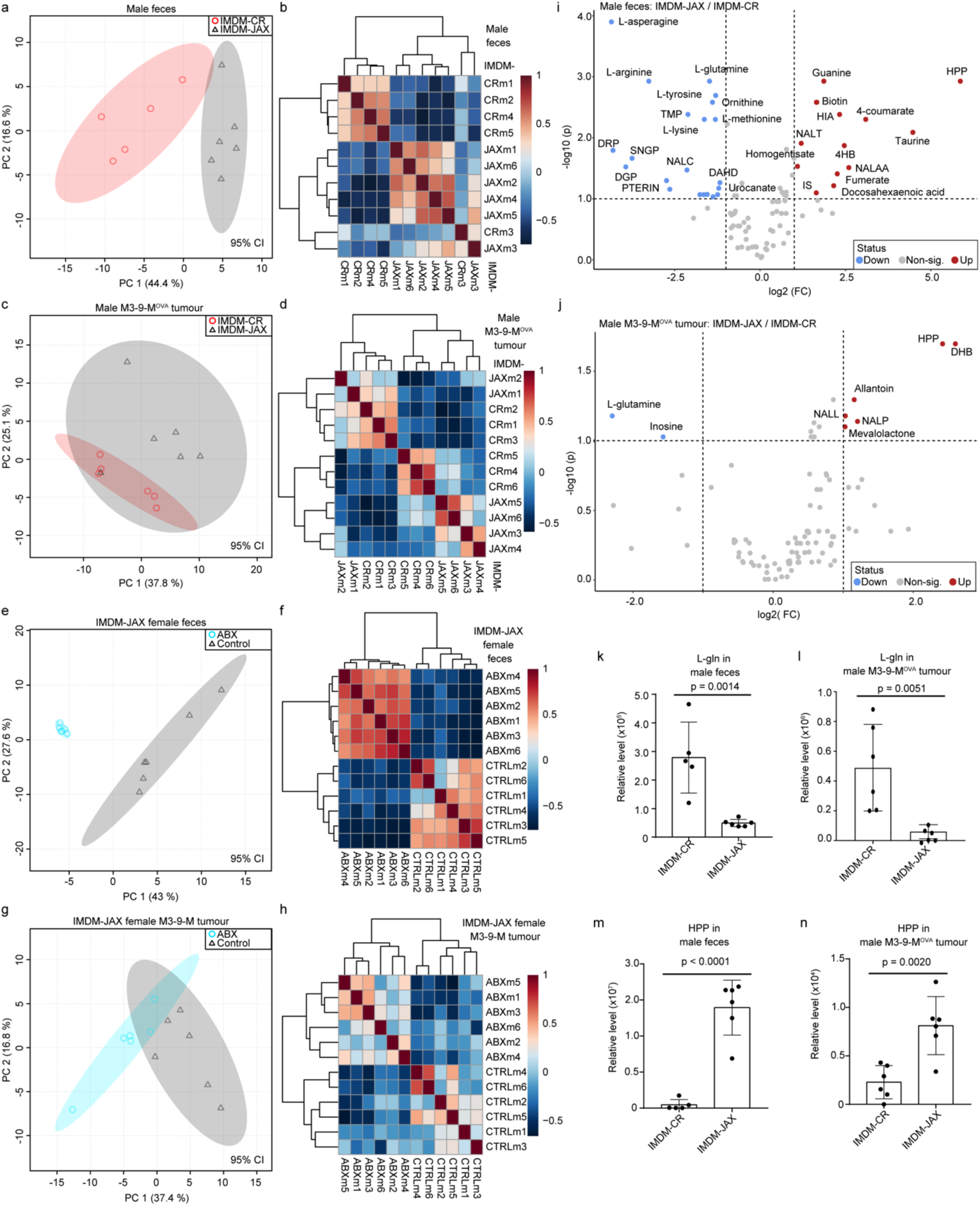
Divergent microbiomes create distinctive metabolomes in the TME. a-b, PC plot and Spearman correlation heatmap of the fecal metabolome of IMDM-CR vs. –JAX male mice (samples pooled from two different experiments, ellipses drawn with 95% confidence interval); c-d, PC plot and Spearman correlation heatmap of orthotopically M3-9-M^OVA^ tumour metabolome of IMDM-CR vs. –JAX male mice (samples pooled from two different experiments, ellipses drawn with 95% confidence interval); e-f, PC plot and Spearman correlation heatmap of the fecal metabolome of IMDM–JAX female mice receiving normal vs. ABX-supplemented water (samples pooled from two different experiments, ellipses drawn with 95% confidence interval); g-h, PC plot and Spearman correlation heatmap of the orthotopically implanted M3-9-M tumour metabolome of IMDM–JAX female mice receiving normal or ABX-supplemented water (samples pooled from two different experiments, ellipses drawn with 95% confidence interval); i-j, volcano plots of the fecal and M3-9-M^OVA^ tumour metabolites of IMDM-CR vs. – JAX male mice, respectively (samples pooled from two different experiments, false discovery rate adjusted); k-n, relative level of select metabolites in feces and M3-9-M^OVA^ tumour of IMDM-CR vs. –JAX male mice (samples pooled from two different experiments, error bars represent standard deviation, unpaired t-tests). The same metabolomic datasets were used to generate different types of graphs for identical experimental groups.

**Extended figure 3:**
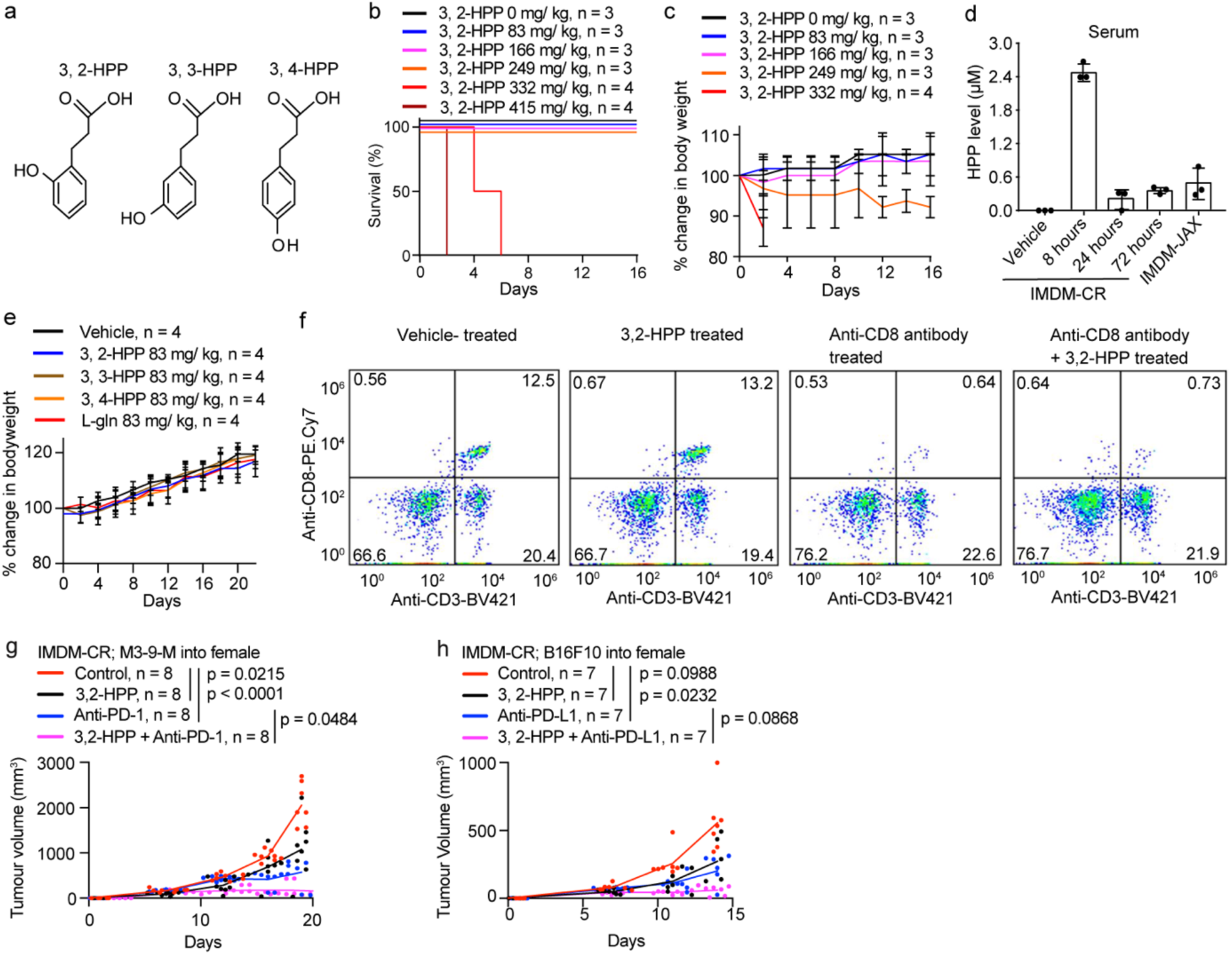
HPP metabolite treatment regimen determination in C57Bl6 mice. a, structure of HPP isomers; b, OS curves for different doses of 3,2-HPP delivered i.p. (single experiment); c, change in mouse body weight with 3,2-HPP treatment (single experiment, error bars represent standard deviation); d, [serum 3,2-HPP] after i.p. treatment with 83 mg/ kg bodyweight of mice (single experiment, error bars represent standard deviation); e, change in mouse body weight with metabolite treatment (single experiment); f, CD8^+^ T cells in mouse tumour 21 days after anti-CD8 neutralizing antibody treatment (2.43 clone; i. p.), by flow cytometry; g-h, tumour growth kinetics in IMDM-CR mice treated with 3,2-HPP (beginning 1 day after cell implantation), anti-PD-1/PD-L1, or both (two combined experiments, two-way ANOVA with Bonferroni tests).

**Extended figure 4:**
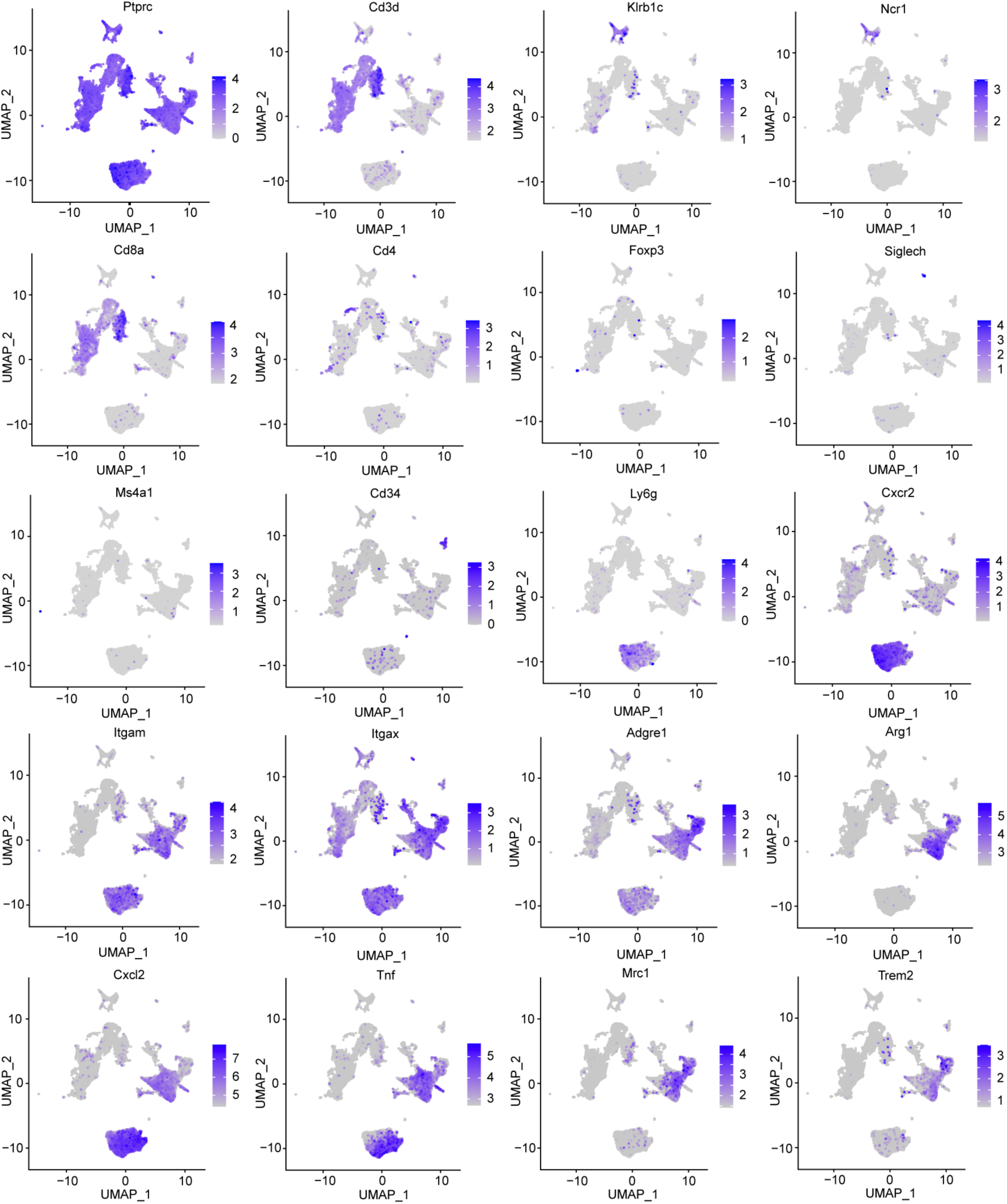
Cell specific marker gene expression used to identify cell clusters obtained through scRNAseq. (cell cluster identity showed in Figure 4a).

**Extended figure 5:**
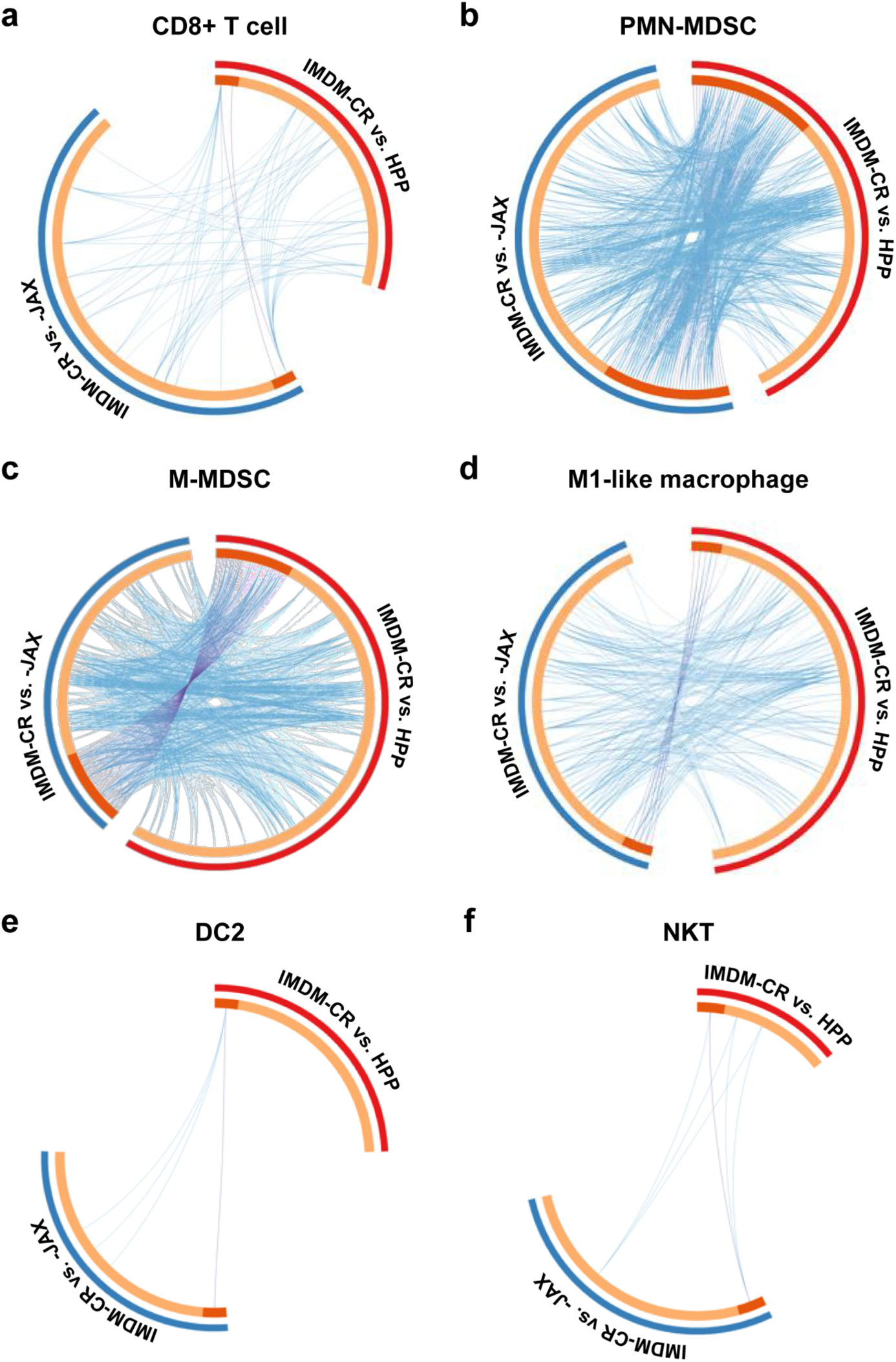
3, 2-HPP metabolite treatment impacts gene expression in different immune cell subsets in the TME. a-f, Circos plots showing overlapping genes that were statistically significantly impacted in the pairwise comparison between IMDM-CR vs. -JAX and IMDM-CR vs. -HPP (Wilcox Rank Sum test).

**Extended figure 6:**
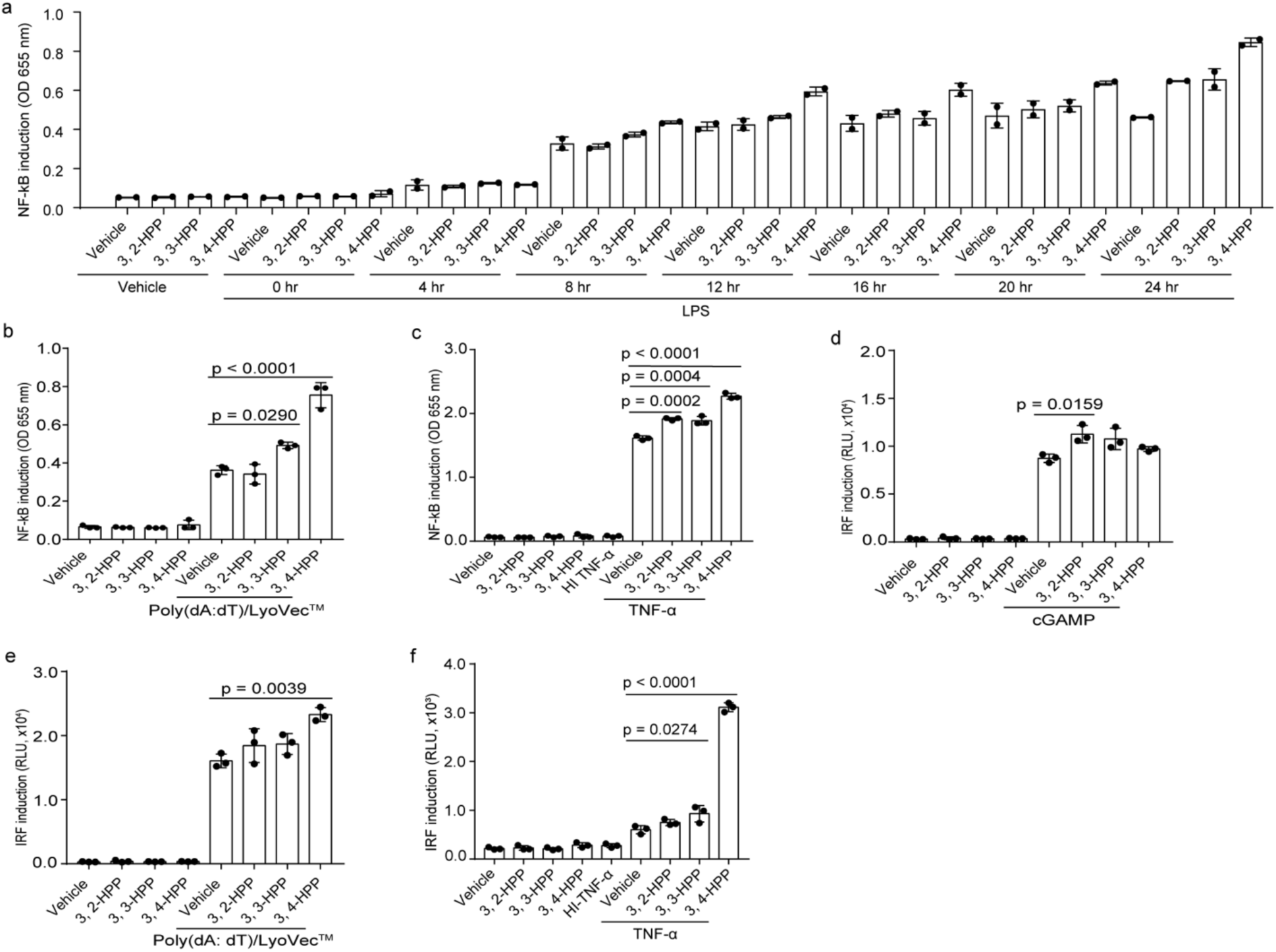
HPP isomers potentiate NF-κB and IRF signalling pathways. a, NF-κB activity in THP1-Dual™ cells treated with LPS in the presence of vehicle or HPP metabolites for 0-24 hours (two combined experiments, error bars represent standard error); b-e, NF-κB or IRF activity in THP1-Dual™ cells treated with different inducers in the presence or absence of HPP isomers for 16 hours (three combined experiments, error bars represent standard error, one-way ANOVA test).

**Extended figure 7:**
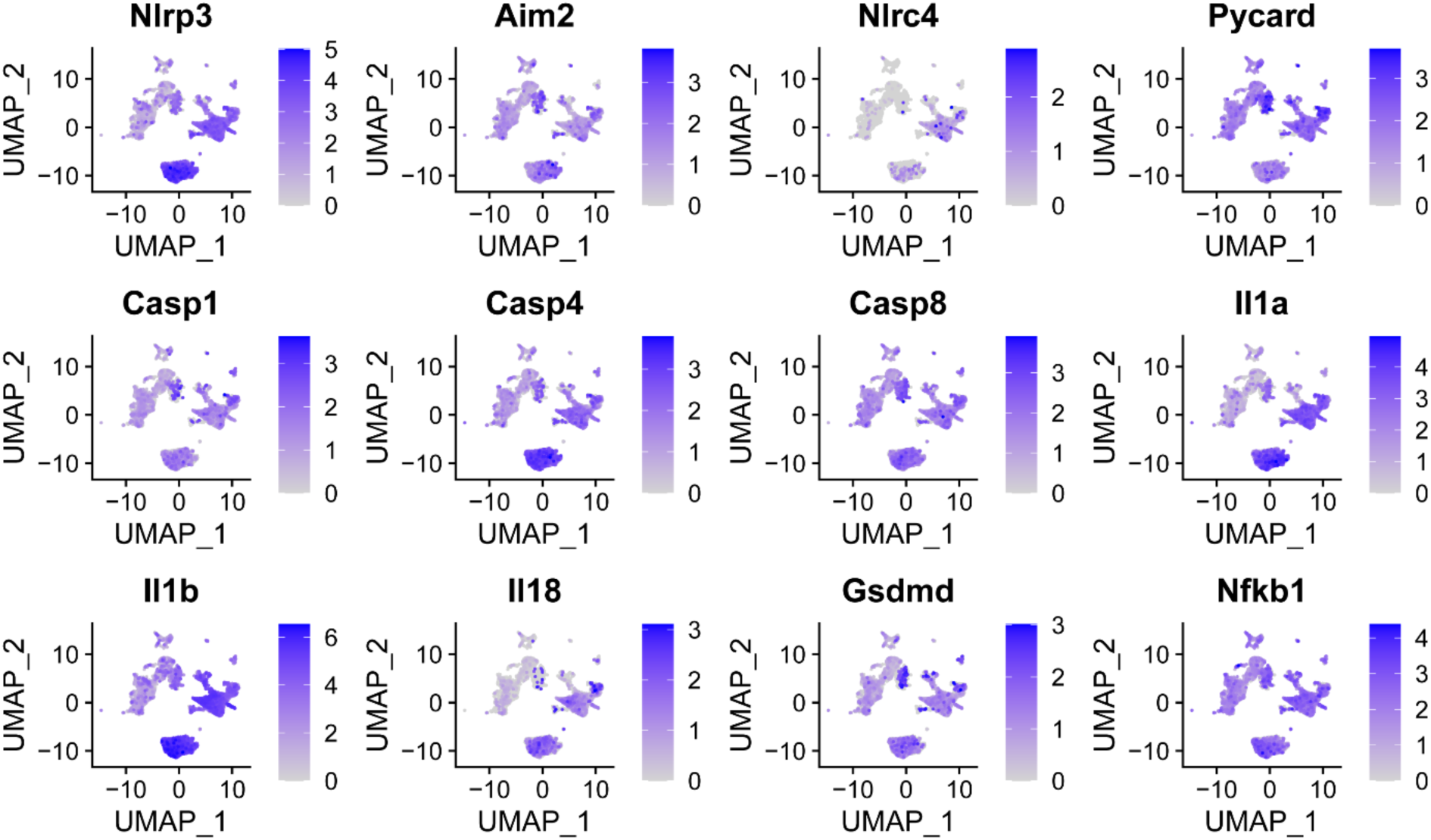
The expression of genes involved in inflammasome pathway identified through scRNAseq. Gene expression in M3-9-M^OVA^ tumour infiltrated leukocytes in IMDM-CR mice receiving 3,2-HPP treatments (cell cluster identity showed in Figure 4a).

**Extended figure 8:**
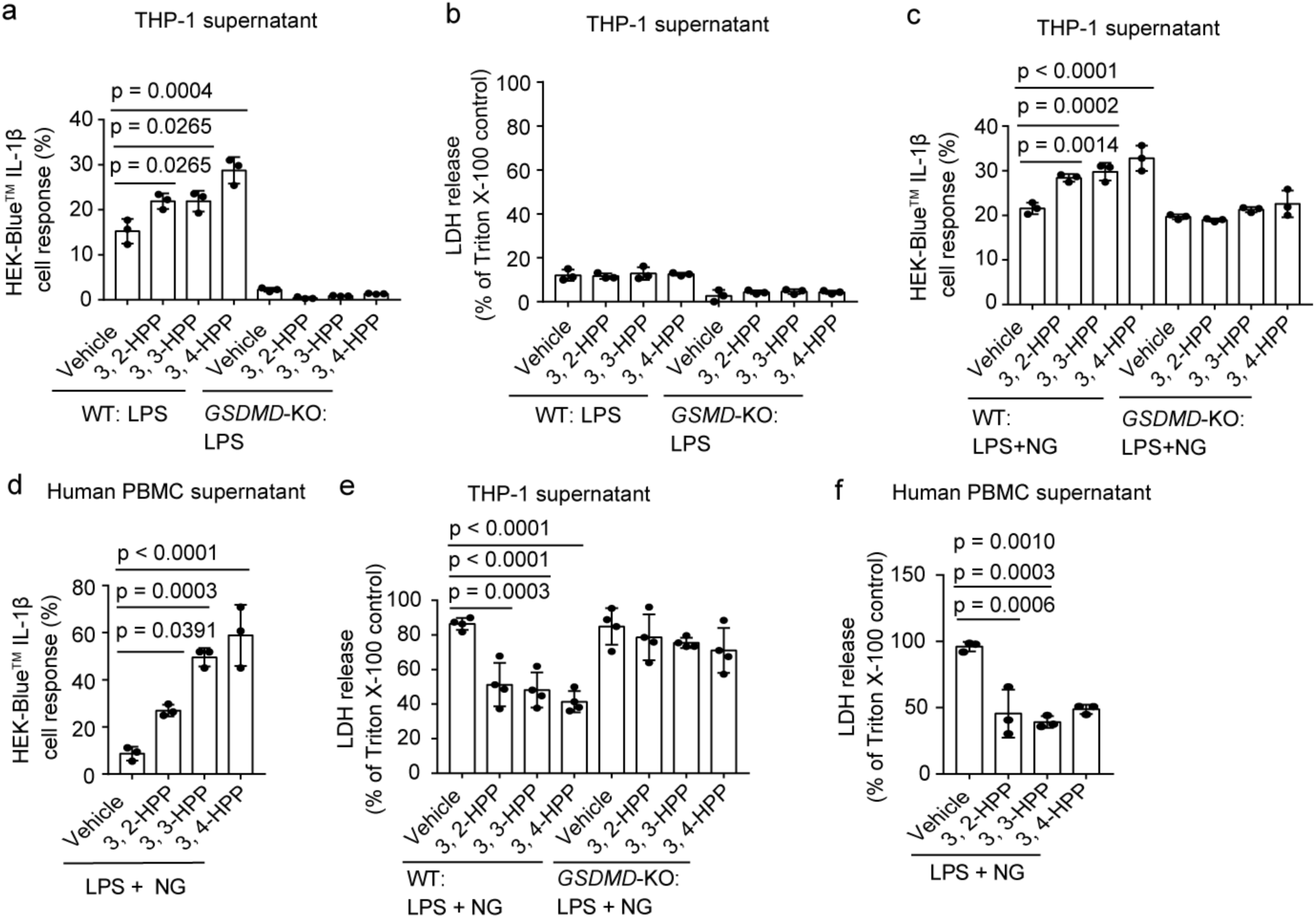
HPP molecules enhance IL-1 receptor signalling while protecting from cell death. a, IL-1β activity in the supernatant of THP1-WT vs. -*GSDMD*-KO cells treated with LPS±HPPs for 16 hours, measured using HEK-Blue™ IL-1β reporter cells (three combined experiments, error bars represent standard error, one-way ANOVA with Bonferroni test); b, LDH release from THP1-WT vs. –*GSDMD*-KO cells treated with LPS±HPP for 16 hours (three combined experiments, one-way ANOVA with Bonferroni test); c-d, IL-1β activity in the supernatant of THP1-WT vs. -*GSDMD*-KO cells © and human PBMCs (d) treated with LPS and NG±HPPs for 16 hours. measured using HEK-Blue™ IL-1β reporter cells (three combined experiments, error bars represent standard error, one-way ANOVA with Bonferroni test); e-f, LDH release from THP1-WT vs. –*GSDMD*-KO cells € and human PBMCs (f) treated with LPS and NG±HPPs for 16 hours (three combined experiments, one-way ANOVA with Bonferroni test).

**Extended figure 9:**
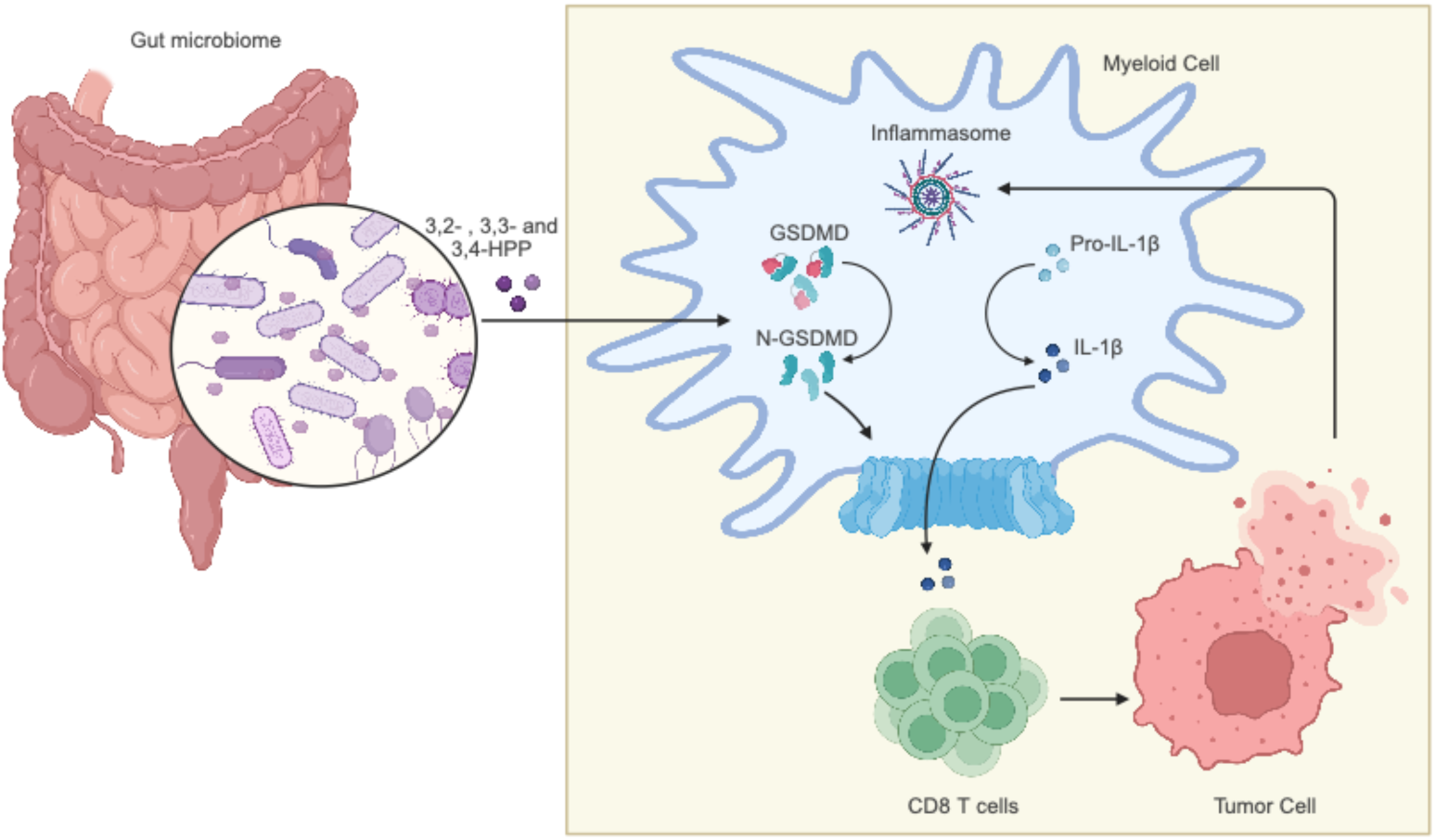
Cartoon representation depicting a proposed mechanisms of how microbiome-derived HPP metabolites modulate antitumour immune response.

**Extended data table 1:**
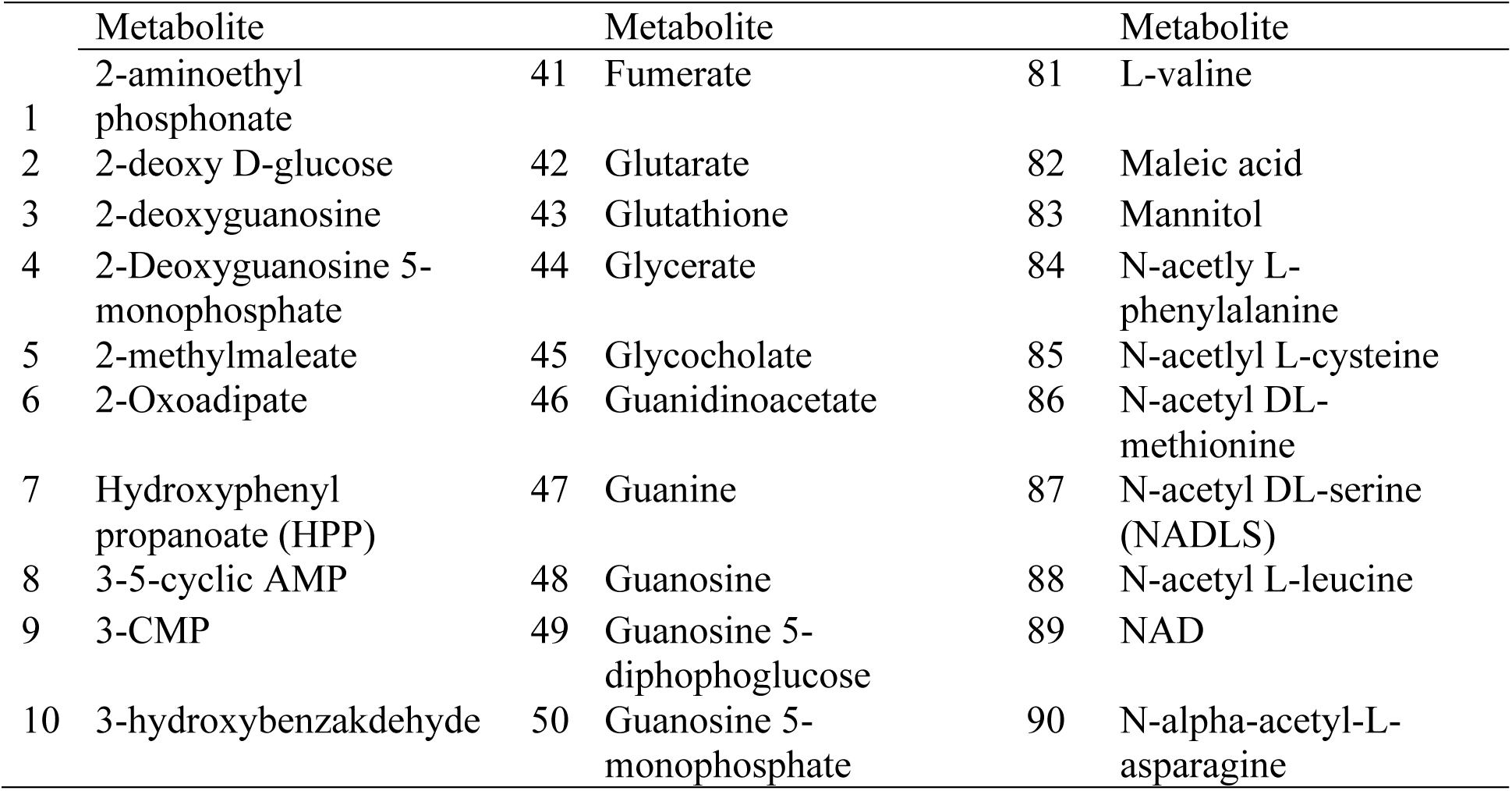
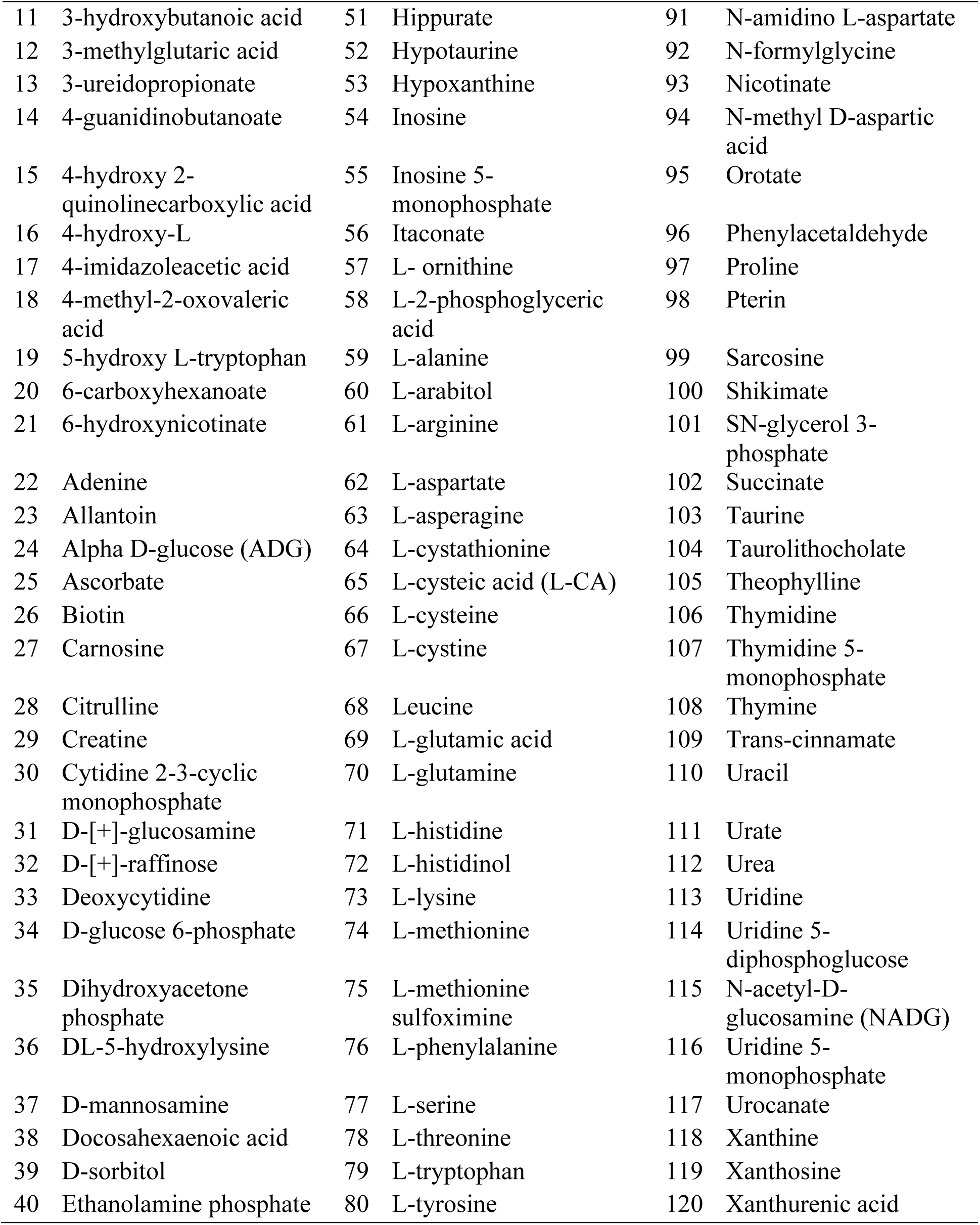
The list of metabolites detected through untargeted UHPLC-MS.

**Extended data table 2:** The list of metabolites enriched in IMDM-CR mice.

| Metabolite name | Gender | Sample type |
| --- | --- | --- |
| L-glutamine | Female, male | Feces, tumour |
| Xanthine | Female, male | Feces, tumour |
| Hippurate | Female, male | Serum, tumour |
| L-asparagine | Female, male | Feces |
| L-arginine | Female, male | Feces |
| L-lysine | Female, male | Feces |
| L-ornithine | Female, male | Feces |
| N-acetyl-dl-serine | Female | Feces |
| 3-methyl-2-oxovaleric acid (MOA) | Female | Feces |
| L-tyrosine | Male | Feces |
| L-glutamic acid | Male | Feces |
| N-acetyl-dl-glutamic acid | Male | Feces |
| Sn-glycerol 3-phosphate | Male | Feces |
| D-glucose 6-phosphate | Male | Feces |
| L-methionine | Male | Feces |
| Chenodeoxycholate | Male | Feces |
| D-ribose 5-phosphate | Male | Feces |
| Urocanate | Male | Feces |
| N-acetyl-l-cysteine | Male | Feces |
| Hypoxanthine | Male | Feces |
| Pterin | Male | Feces |
| Alpha-hydroxyisobutyric acid | Male | Feces |
| 3-ureidopropionate | Male | Feces |
| Xanthosine | Male | Feces |
| Salicylate | Female | Serum |
| 2-quinolinecarboxylic acid | Female | Tumour |
| Mono-ethyl malonate | Female | Tumour |

**Extended data table 3:**
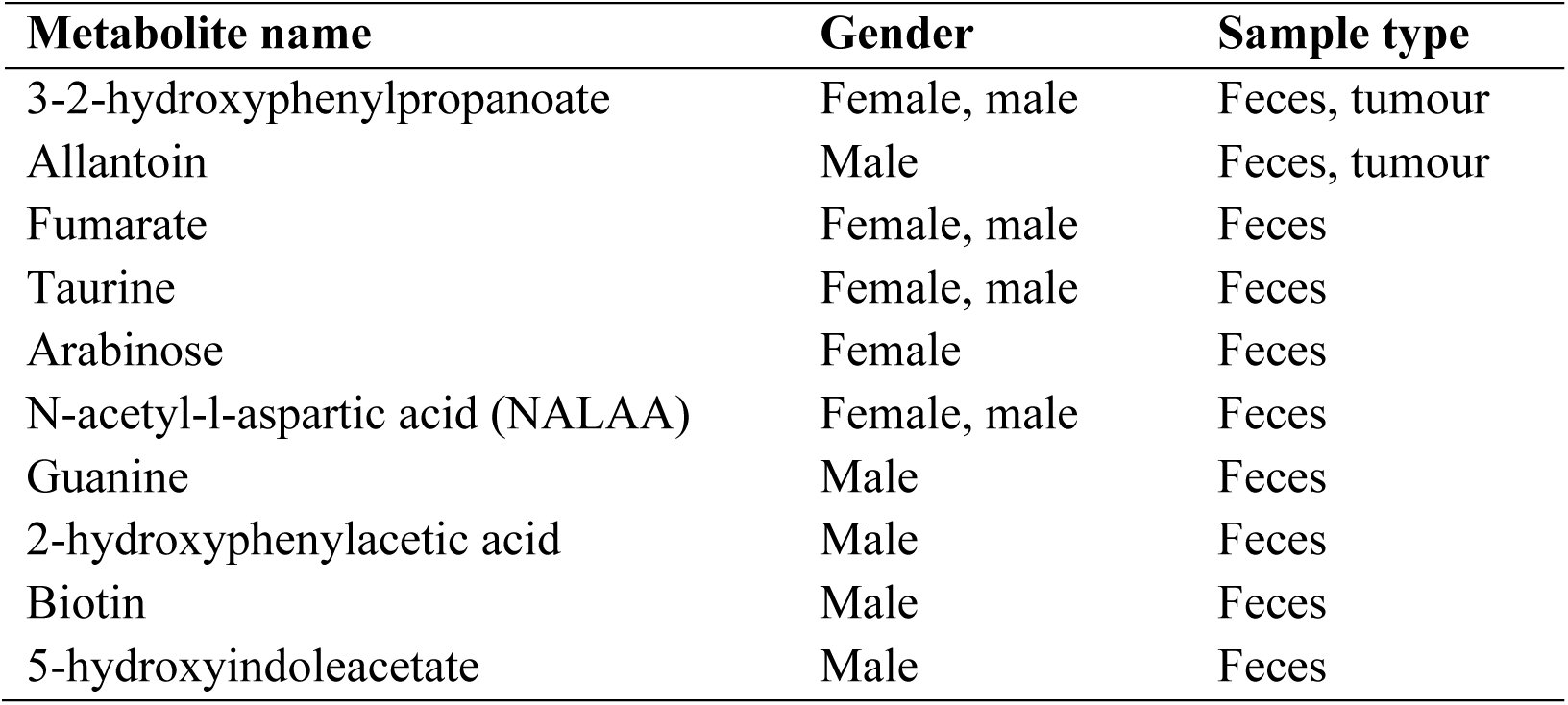
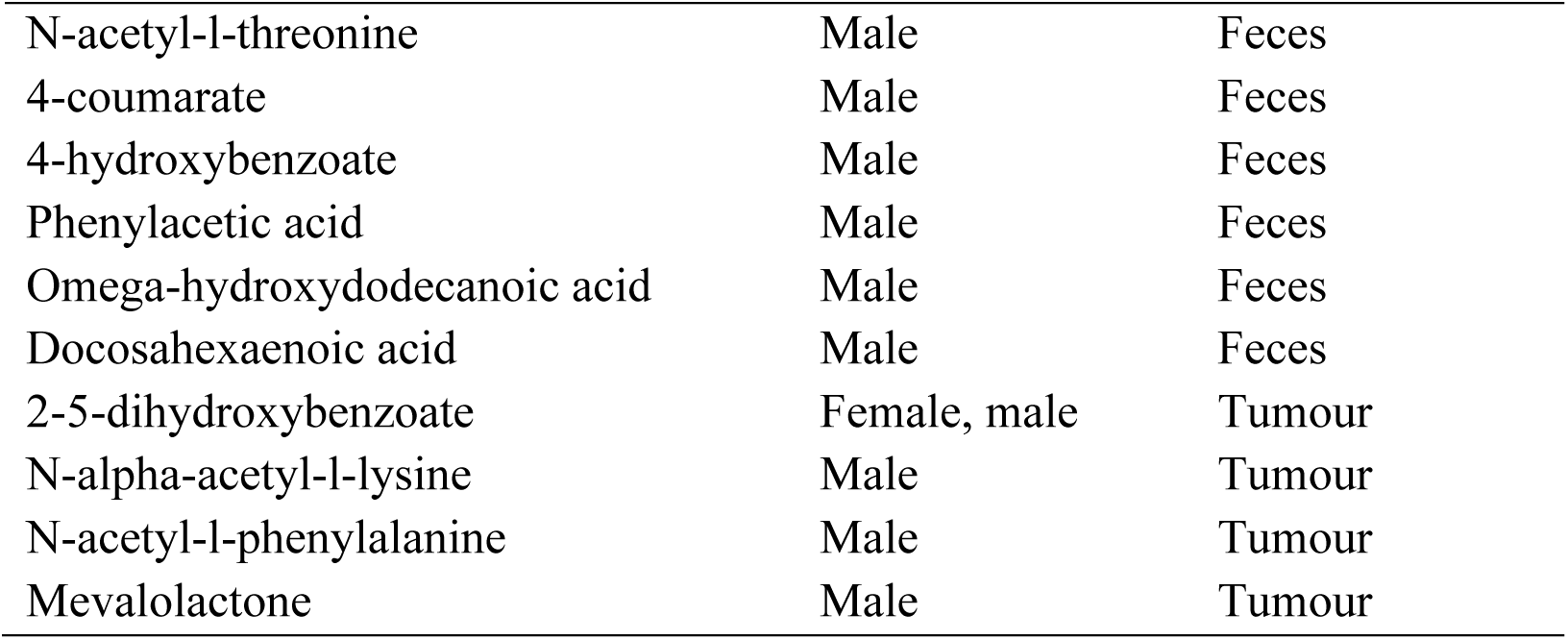
The list of metabolites enriched in IMDM-JAX mice.

**Extended data table 4:** Members of the microbiome positively associated with HPP metabolite production. The machine learning tool Melonnpan identifies members of the microbiome that produce HPP metabolite (Spearman = 0.9860, p-value < 0.0001, q-value < 0.0001 and model size = 149).

| ID | Name | Coefficient |
| --- | --- | --- |
| Intercept | NA | 0.2623 |
| ASV_0454 | *Muribaculaceae* | 0.0025 |
| ASV_0202 | *Muribaculaceae* | 0.0024 |
| ASV_0216 | *Muribaculaceae* | 0.0021 |
| ASV_0334 | *Muribaculaceae* | 0.0021 |
| ASV_1419 | *Erysipelotrichaceae* | 0.0019 |
| ASV_1245 | *Erysipelatoclostridiaceae* | 0.0017 |
| ASV_0978 | *Erysipelotrichaceae* | 0.0017 |
| ASV_0008 | *Muribaculaceae* | 0.0017 |
| ASV_0337 | *Clostridium_sensu_stricto_1* | 0.0017 |
| ASV_0009 | *Muribaculaceae* | 0.0015 |
| ASV_0310 | *Muribaculaceae* | 0.0015 |
| ASV_2014 | *Muribaculaceae* | 0.0014 |
| ASV_0294 | *Clostridium_sensu_stricto_1* | 0.0014 |
| ASV_1689 | *Muribaculaceae* | 0.0014 |
| ASV_0422 | *Clostridium_sensu_stricto_1* | 0.0013 |
| ASV_0973 | *Muribaculaceae* | 0.0013 |
| ASV_1381 | *Parasutterella* | 0.0013 |
| ASV_2274 | *Muribaculaceae* | 0.0013 |
| ASV_0025 | *Muribaculaceae* | 0.0012 |
| ASV_0105 | *Parasutterella* | 0.0012 |
| ASV_0019 | *Muribaculaceae* | 0.0012 |
| ASV_1850 | *Adlercreutzia* | 0.0011 |
| ASV_1269 | *Muribaculaceae* | 0.0011 |
| ASV_0863 | *Clostridia_vadinBB60_group* | 0.0011 |
| ASV_0284 | *Muribaculaceae* | 0.0011 |
| ASV_0033 | *Muribaculaceae* | 0.0011 |
| ASV_1019 | *Erysipelatoclostridiaceae* | 0.0010 |
| ASV_0606 | *Marvinbryantia* | 0.0010 |
| ASV_0166 | *Muribaculaceae* | 0.0010 |
| ASV_0480 | *Clostridium_sensu_stricto_1* | 0.0010 |
| ASV_0852 | *Alistipes* | 0.0010 |
| ASV_0316 | *Muribaculaceae* | 0.0009 |
| ASV_0195 | *Adlercreutzia* | 0.0009 |
| ASV_0552 | *Parvibacter* | 0.0009 |
| ASV_0532 | *Adlercreutzia* | 0.0009 |
| ASV_0432 | *Eggerthellaceae* | 0.0009 |
| ASV_1794 | *Clostridia_vadinBB60_group* | 0.0009 |
| ASV_1199 | *Muribaculaceae* | 0.0009 |
| ASV_0050 | *Muribaculaceae* | 0.0009 |
| ASV_0286 | *Eggerthellaceae* | 0.0008 |
| ASV_0880 | *Muribaculaceae* | 0.0008 |
| ASV_1204 | *Merdibacter* | 0.0008 |
| ASV_0593 | *Clostridium_sensu_stricto_1* | 0.0008 |
| ASV_0326 | *Adlercreutzia* | 0.0008 |
| ASV_1344 | *Lachnospiraceae* | 0.0008 |
| ASV_0044 | *Muribaculaceae* | 0.0007 |
| ASV_0024 | *Coriobacteriaceae_UCG-002* | 0.0007 |
| ASV_0338 | *Colidextribacter* | 0.0006 |
| ASV_0719 | *Adlercreutzia* | 0.0006 |
| ASV_0637 | *Eggerthellaceae* | 0.0006 |
| ASV_0589 | *Muribaculaceae* | 0.0006 |
| ASV_0787 | *Marvinbryantia* | 0.0006 |
| ASV_1798 | *Lactobacillus* | 0.0006 |
| ASV_0113 | *Muribaculaceae* | 0.0005 |
| ASV_0158 | *Parasutterella* | 0.0005 |
| ASV_0488 | *Muribaculaceae* | 0.0005 |
| ASV_0735 | *Parvibacter* | 0.0005 |
| ASV_0254 | *Monoglobus* | 0.0005 |
| ASV_0261 | *Muribaculaceae* | 0.0004 |
| ASV_0281 | *Muribaculaceae* | 0.0004 |
| ASV_1291 | *Muribaculaceae* | 0.0004 |
| ASV_0069 | *Muribaculaceae* | 0.0004 |
| ASV_0012 | *Muribaculaceae* | 0.0004 |
| ASV_0390 | *Muribaculaceae* | 0.0003 |
| ASV_0465 | *Clostridia_UCG-014* | 0.0003 |
| ASV_0236 | *Marvinbryantia* | 0.0003 |
| ASV_0032 | *Coriobacteriaceae_UCG-002* | 0.0003 |
| ASV_0020 | *Muribaculaceae* | 0.0003 |
| ASV_0102 | *Muribaculaceae* | 0.0003 |
| ASV_1690 | *Muribaculaceae* | 0.0003 |
| ASV_0160 | *Monoglobus* | 0.0002 |
| ASV_1039 | *Muribaculaceae* | 0.0002 |
| ASV_0641 | *Clostridia_UCG-014* | 0.0002 |
| ASV_1119 | *Clostridia_UCG-014* | 0.0002 |
| ASV_0081 | *Muribaculaceae* | 0.0001 |
| ASV_0180 | *Muribaculaceae* | 0.0001 |
| ASV_0034 | *Muribaculaceae* | 0.0001 |

**Extended data table 5:**
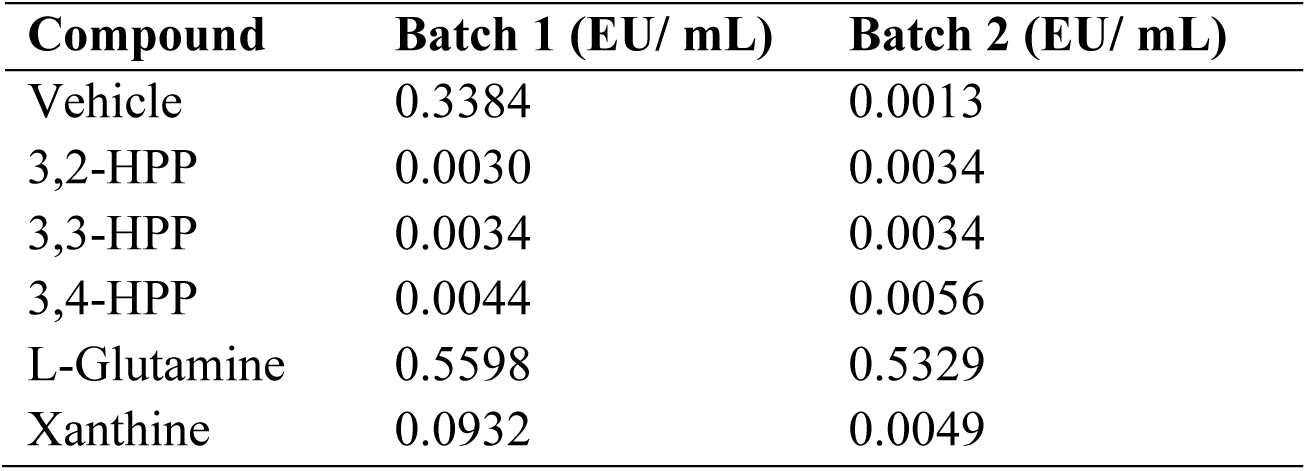
LAL test determines endotoxin level of the metabolite solutions used in the present study.

| Compound | Batch 1 (EU/ mL) | Batch 2 (EU/ mL) |
| --- | --- | --- |
| Vehicle | 0.3384 | 0.0013 |
| 3,2-HPP | 0.0030 | 0.0034 |
| 3,3-HPP | 0.0034 | 0.0034 |
| 3,4-HPP | 0.0044 | 0.0056 |
| L-Glutamine | 0.5598 | 0.5329 |
| Xanthine | 0.0932 | 0.0049 |

**Extended data table 6:**
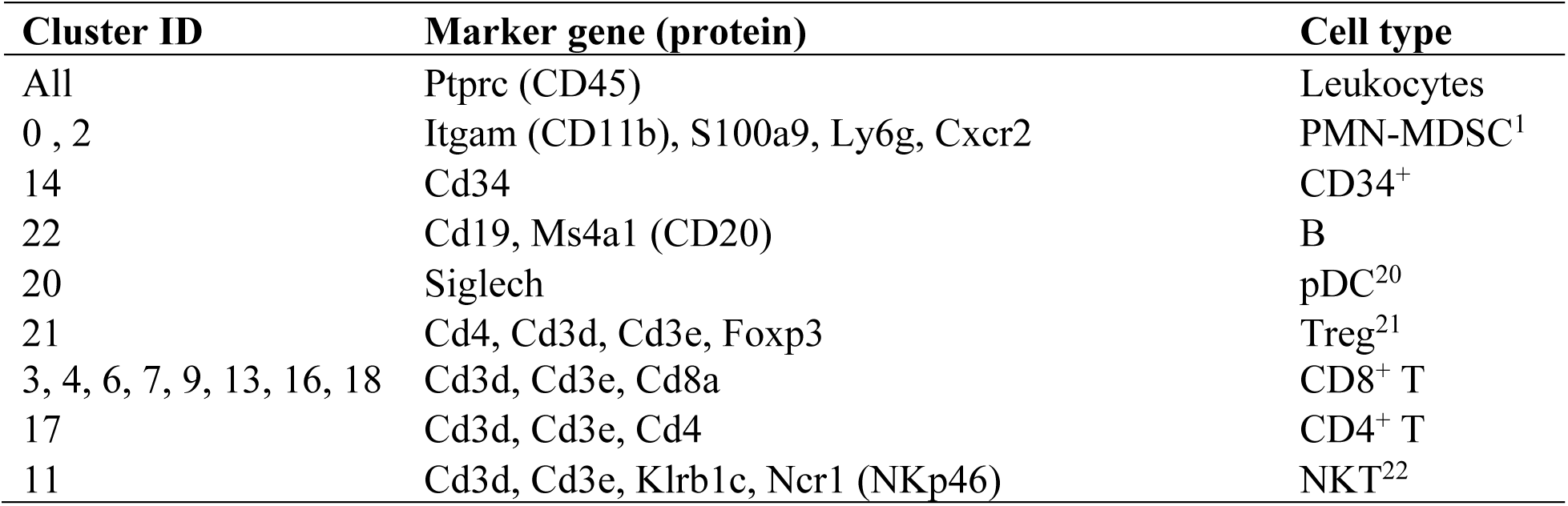
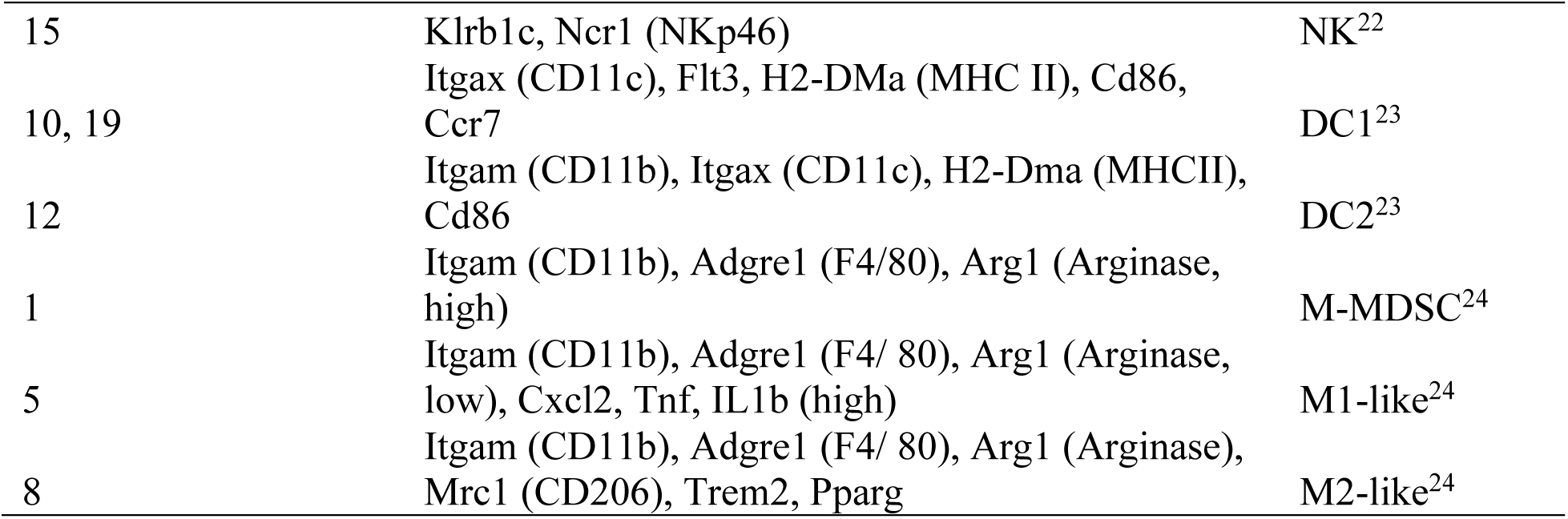
Cell identification marker genes used to annotate the unsupervised clusters obtained from scRNAseq.

**Extended data table 7:**
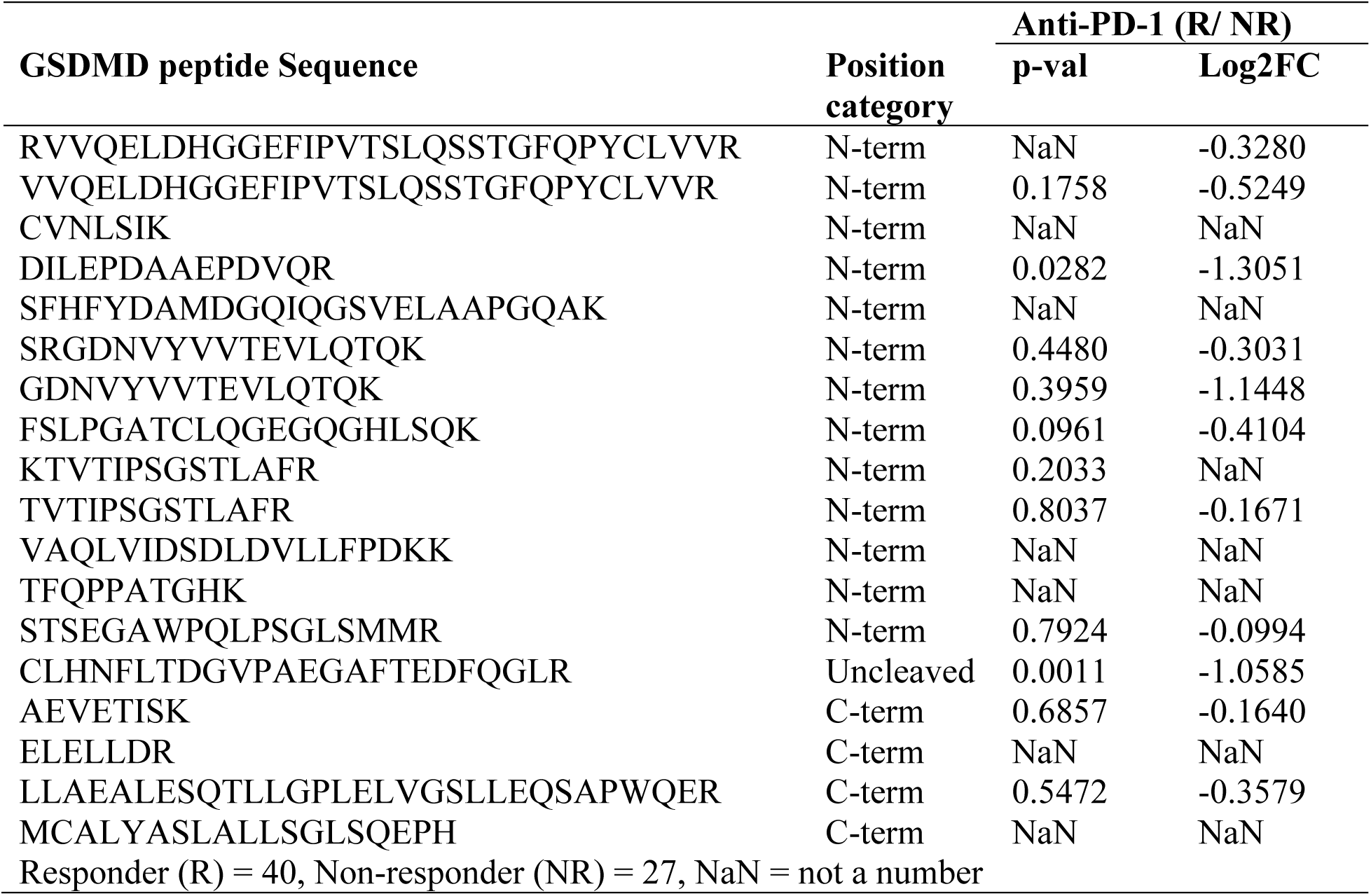
GDSMD peptide quantification from a publicly available proteomic dataset of advanced stage melanoma patients. The proteomic data was obtained from clinical samples of stage IV melanoma patients undergoing anti-PD-1 immunotherapy^25^.

## Notes

### Competing Interest Statement

DJM and SS are cofounders of Oncobiotix Inc.

